## Supplementary material 2, and will be used for the link to the file on the preprint site. for "Species and habitat specific changes in bird activity in an urban environment during Covid 19 lockdown"

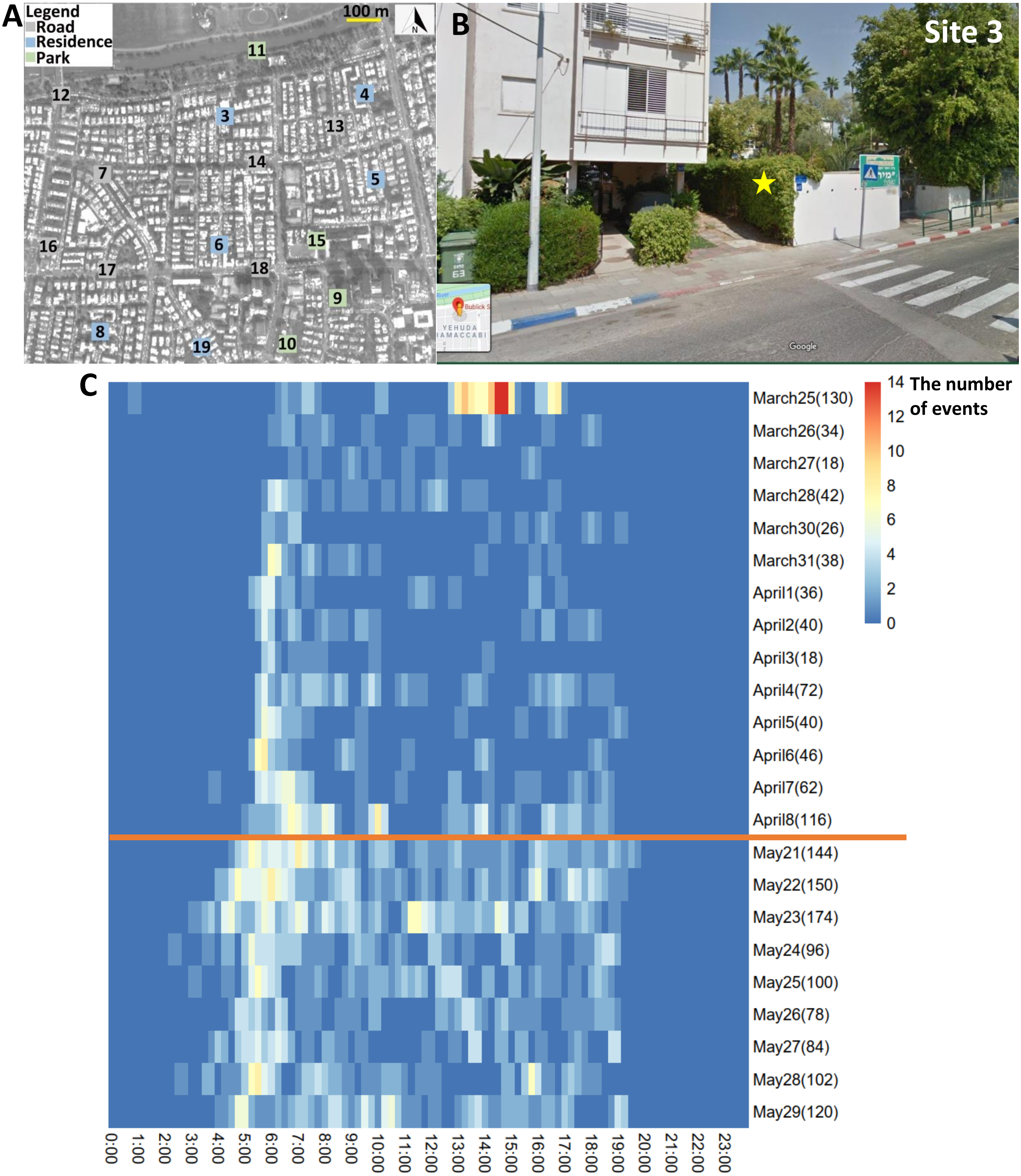


**Figure S1.** (A) Study area and (B) recording site 3. The yellow star refers to the audiomoth's location. (C) Heatmap indicating the activity of *Corvus corone cornix* along the day. The x-axis refers to the time of day. The y-axis is the date. The numbers in parentheses for dates represent the total number of events detected during the day. The orange line separates lockdown from no lockdown periods.

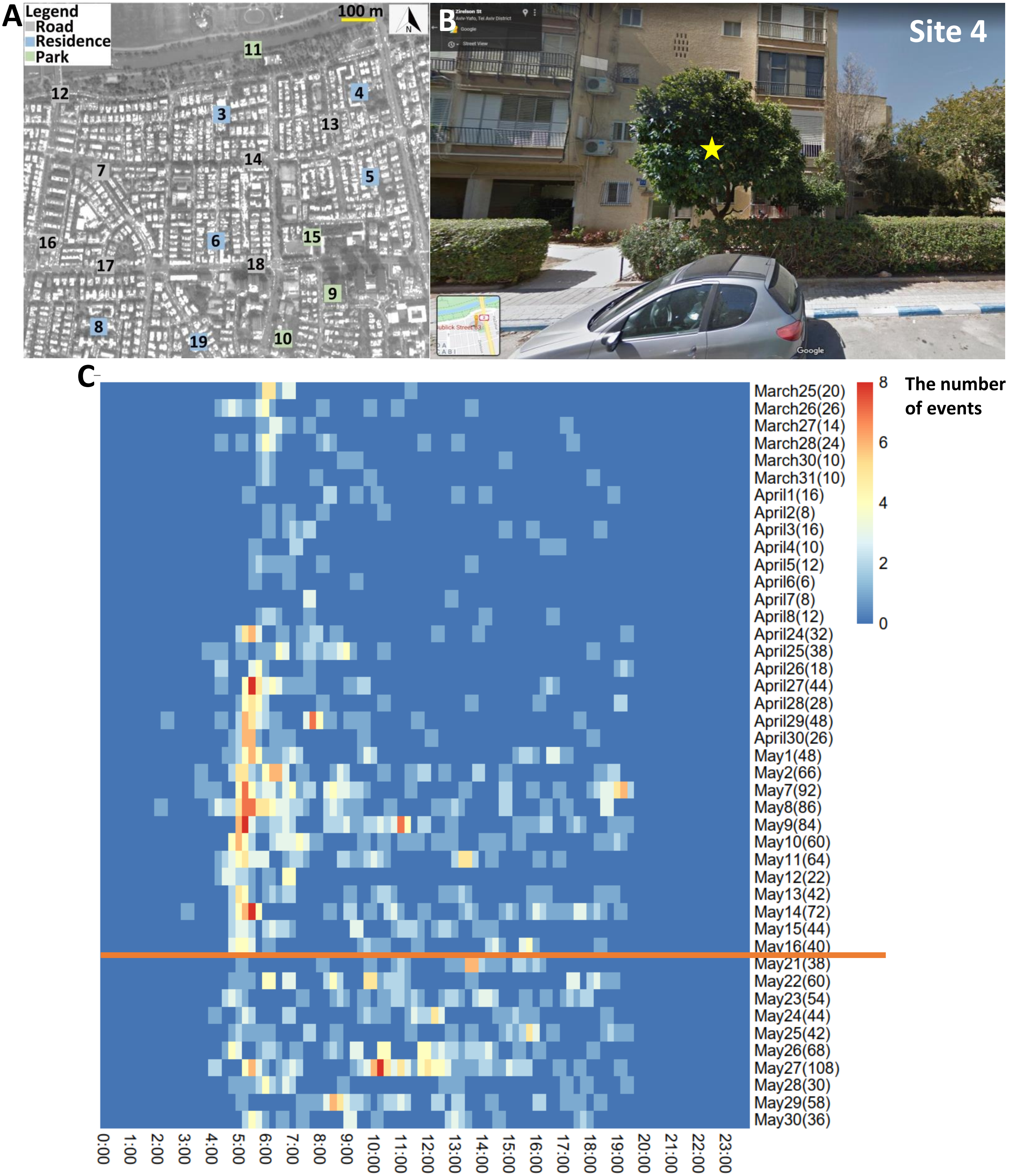


**Figure S2.** (A) Study area and (B) recording site 4. The yellow star refers to the audiomoth's location. (C) Heatmap indicating the activity of *Corvus corone cornix* along the day. The x-axis refers to the time of day. The y-axis is the date. The numbers in parentheses for dates represent the total number of events detected during the day. The orange line separates lockdown from no lockdown periods.

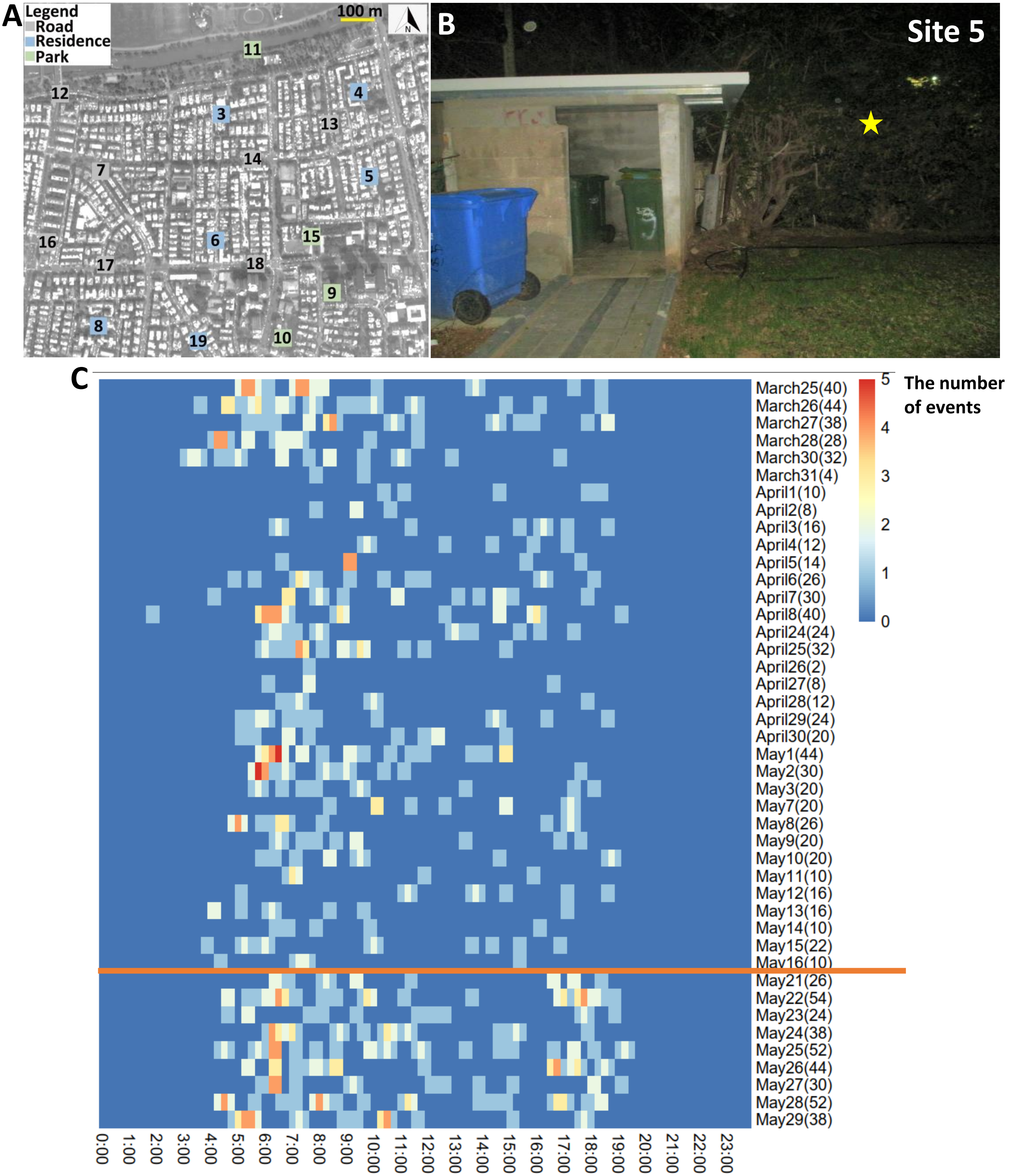


**Figure S3.** (A) Study area and (B) recording site 5. The yellow star refers to the audiomoth's location. (C) Heatmap indicating the activity of *Corvus corone cornix* along the day. The x-axis refers to the time of day. The y-axis is the date. The numbers in parentheses for dates represent the total number of events detected during the day. The orange line separates lockdown from no lockdown periods.

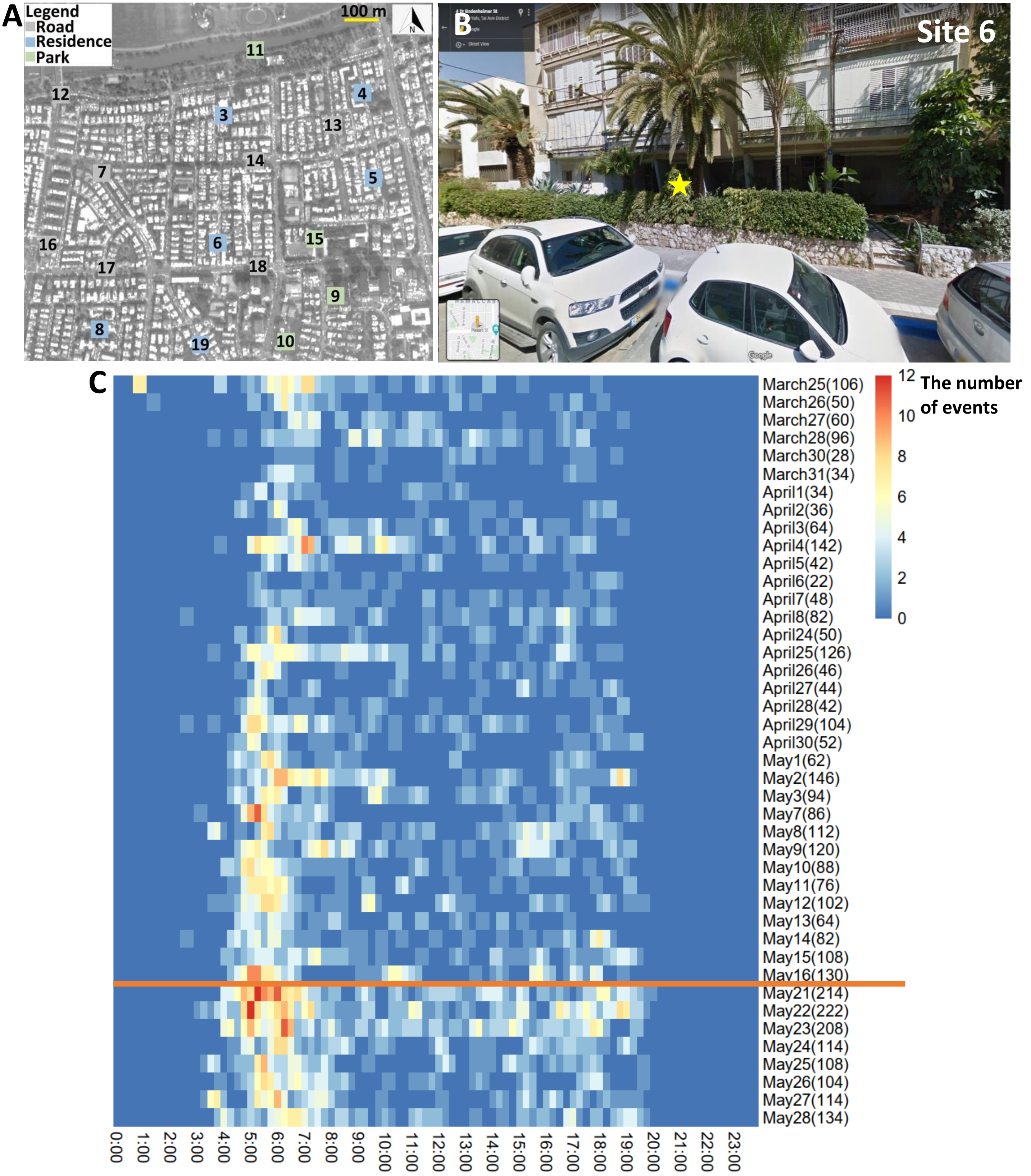


**Figure S4.** (A) Study area and (B) recording site 6. The yellow star refers to the audiomoth's location. (C) Heatmap indicating the activity of *Corvus corone cornix* along the day. The x-axis refers to the time of day. The y-axis is the date. The numbers in parentheses for dates represent the total number of events detected during the day. The orange line separates lockdown from no lockdown periods.

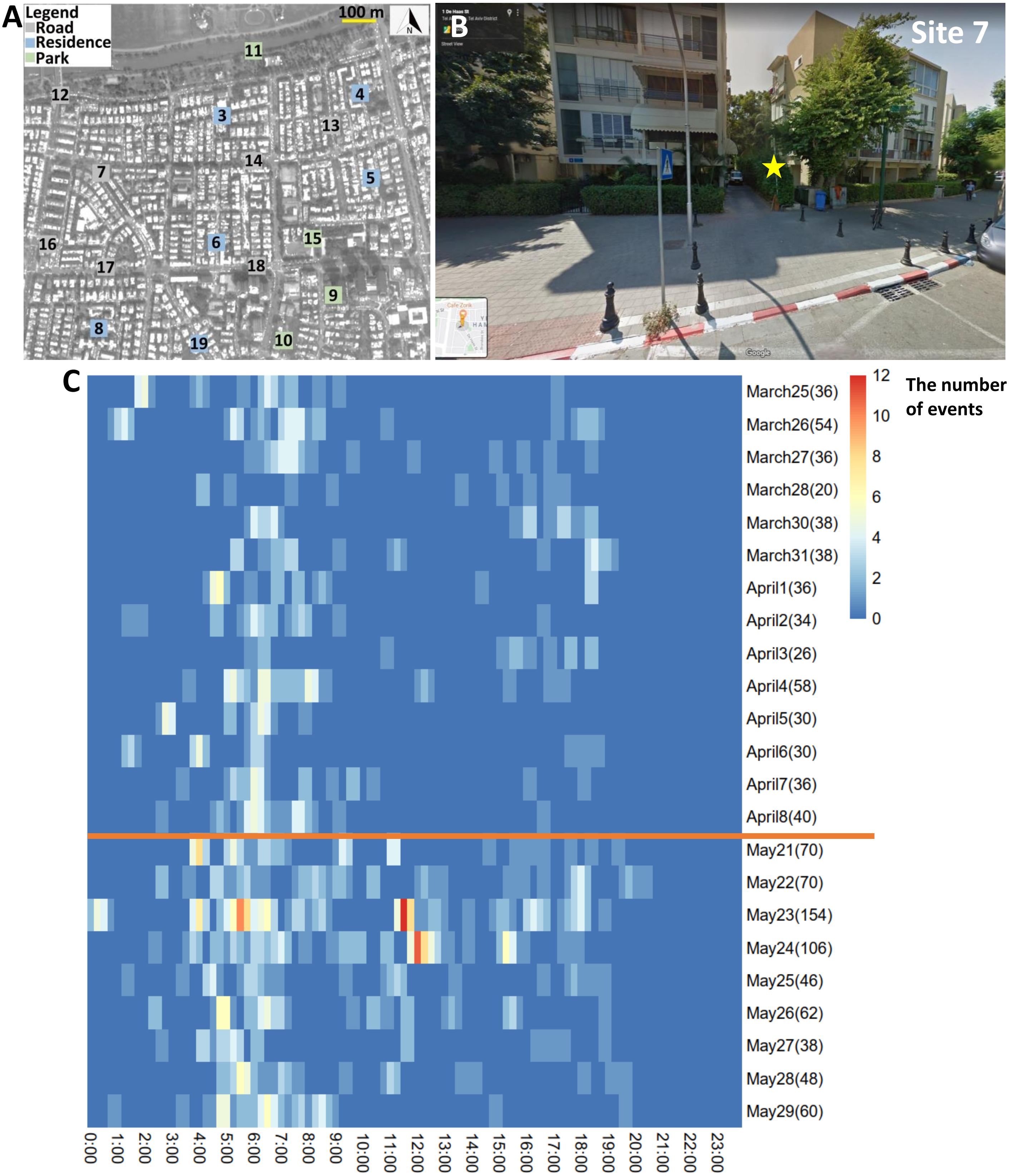


**Figure S5.** (A) Study area and (B) recording site 7. The yellow star refers to the audiomoth's location. (C) Heatmap indicating the activity of *Corvus corone cornix* along the day. The x-axis refers to the time of day. The y-axis is the date. The numbers in parentheses for dates represent the total number of events detected during the day. The orange line separates lockdown from no lockdown periods.

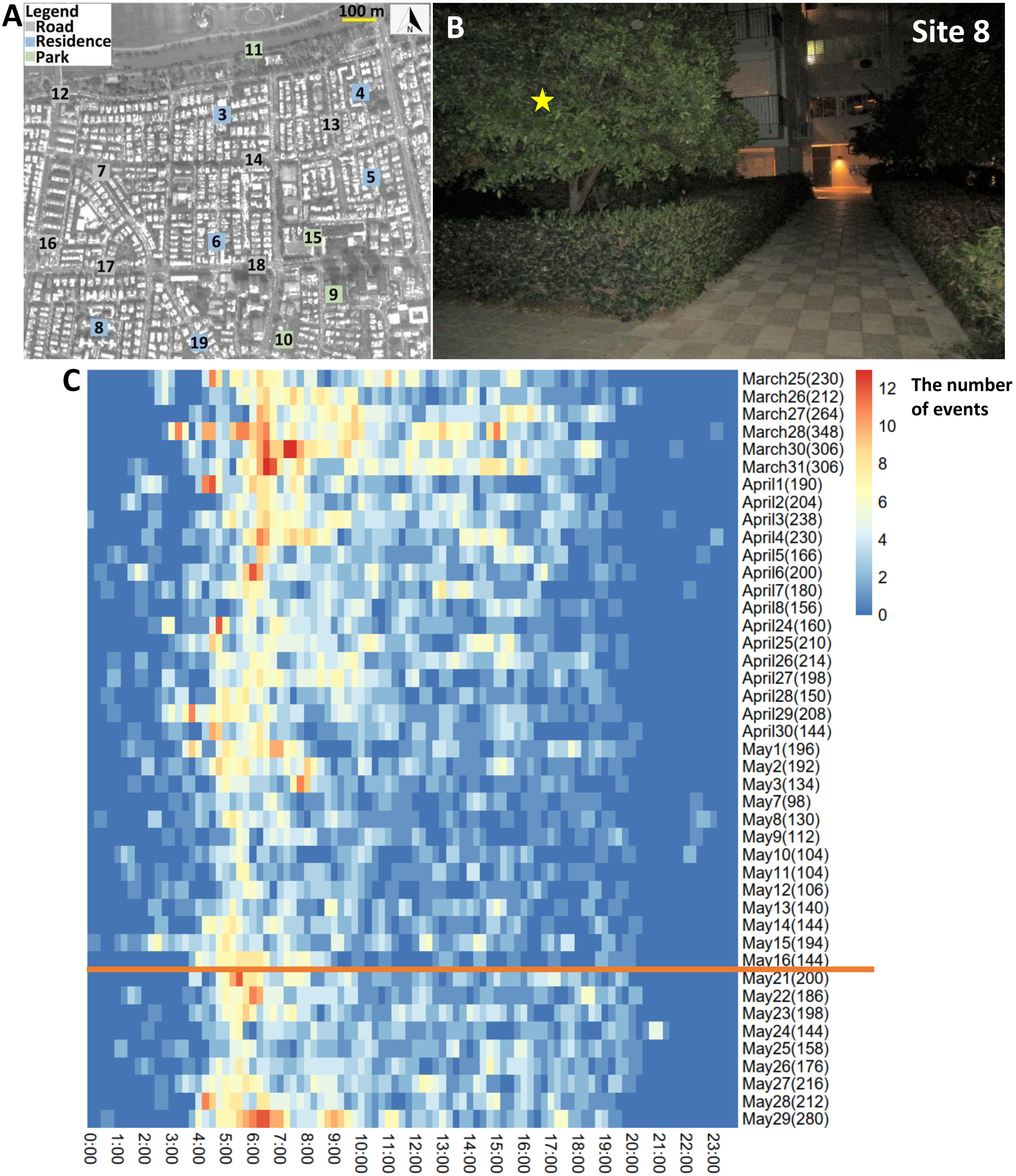


**Figure S6.** (A) Study area and (B) recording site 8. The yellow star refers to the audiomoth's location. (C) Heatmap indicating the activity of *Corvus corone cornix* along the day. The x-axis refers to the time of day. The y-axis is the date. The numbers in parentheses for dates represent the total number of events detected during the day. The orange line separates lockdown from no lockdown periods.

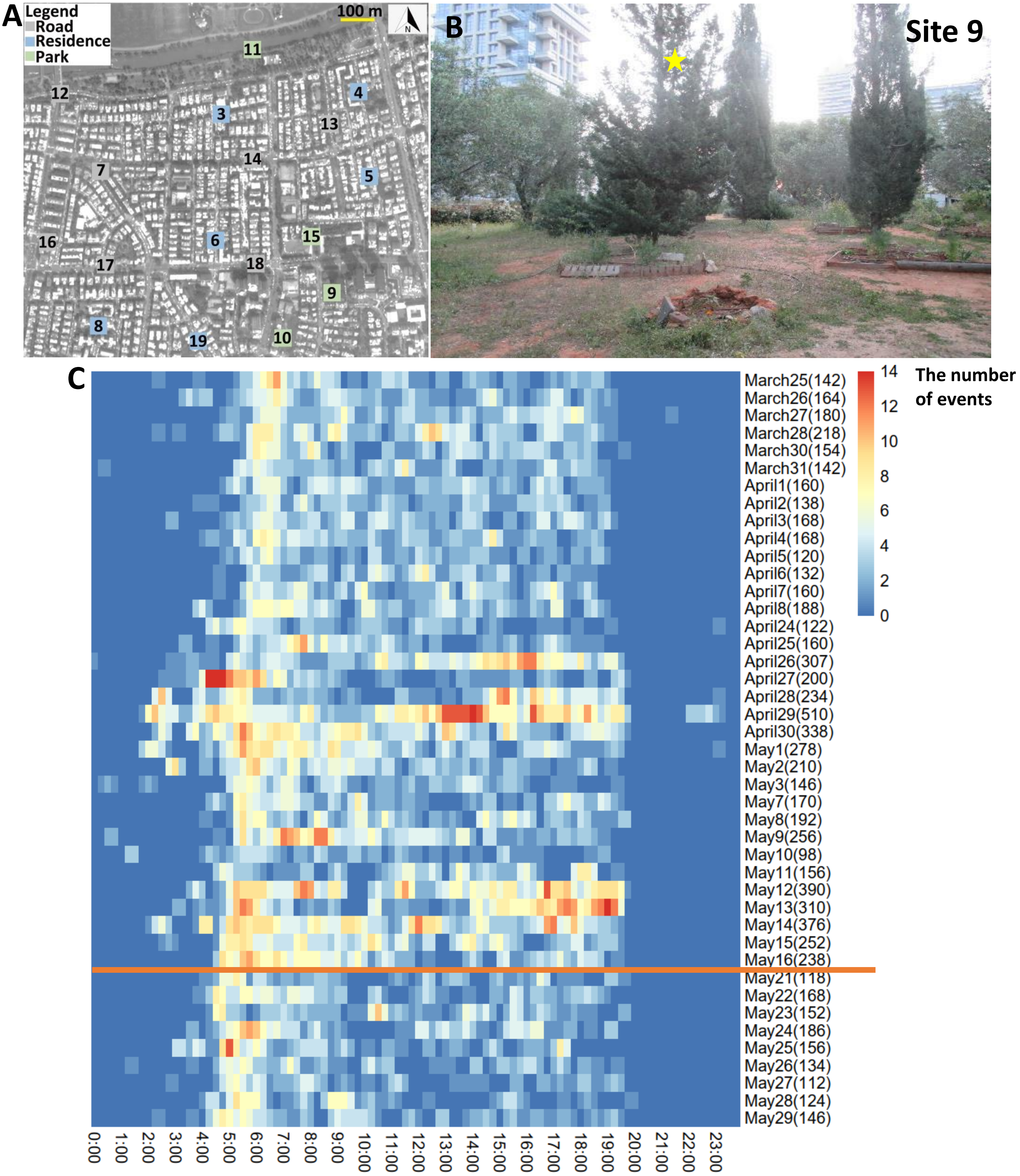


**Figure S7.** (A) Study area and (B) recording site 9. The yellow star refers to the audiomoth's location. (C) Heatmap indicating the activity of *Corvus corone cornix* along the day. The x-axis refers to the time of day. The y-axis is the date. The numbers in parentheses for dates represent the total number of events detected during the day. The orange line separates lockdown from no lockdown periods.

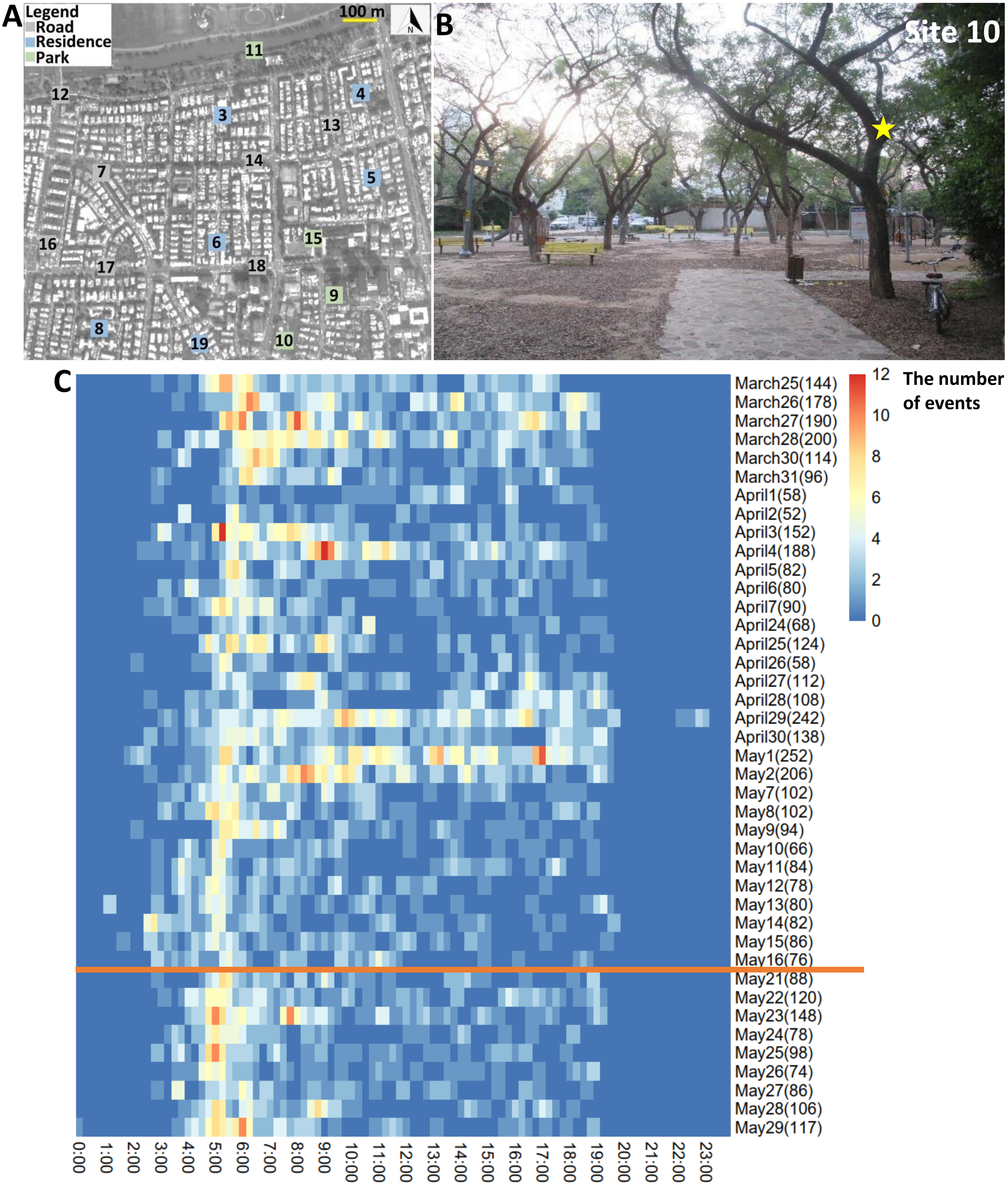


**Figure S8.** (A) Study area and (B) recording site 10. The yellow star refers to the audiomoth's location. (C) Heatmap indicating the activity of *Corvus corone cornix* along the day. The x-axis refers to the time of day. The y-axis is the date. The numbers in parentheses for dates represent the total number of events detected during the day. The orange line separates lockdown from no lockdown periods.

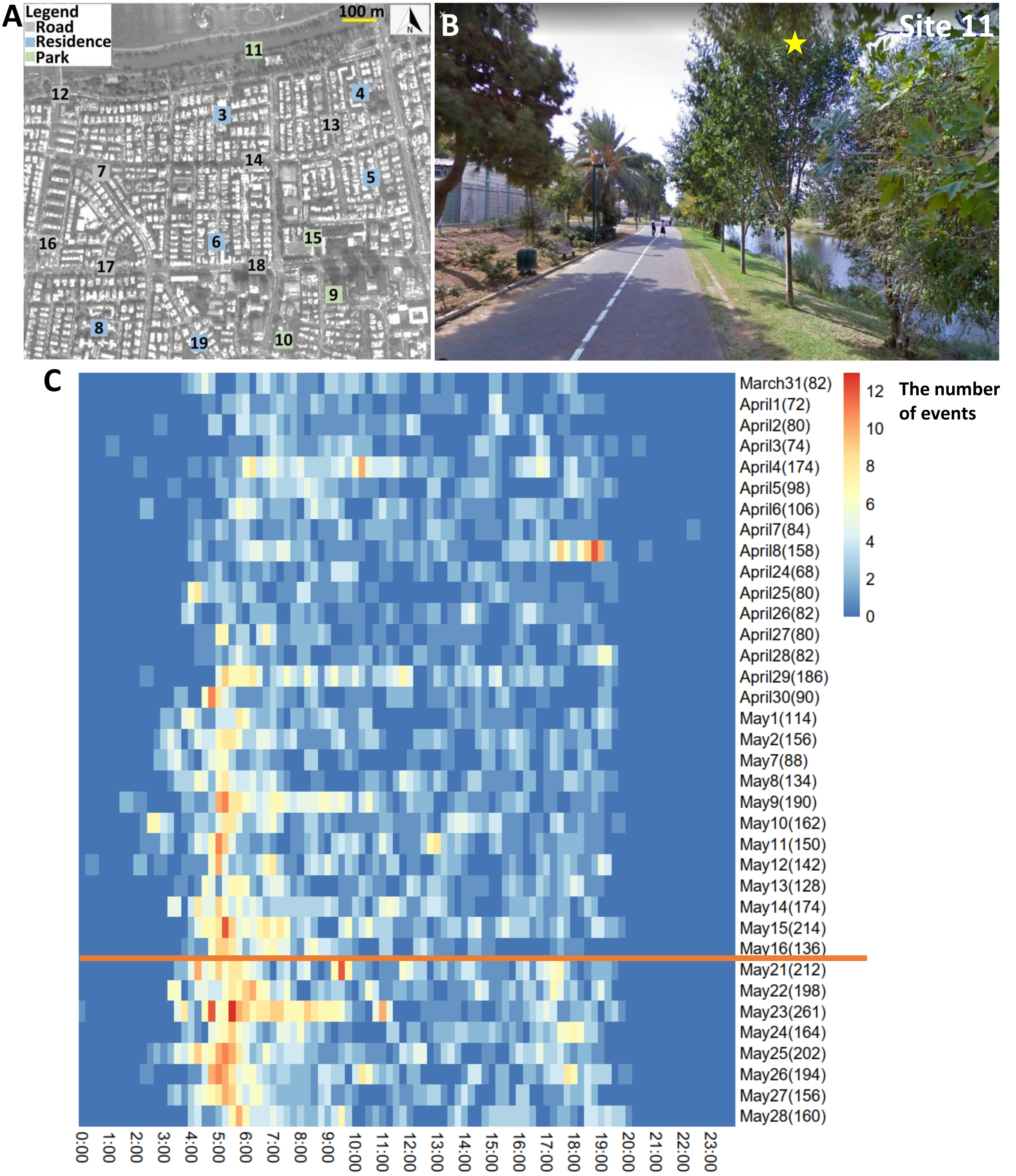


**Figure S9.** (A) Study area and (B) recording site 11. The yellow star refers to the audiomoth's location. (C) Heatmap indicating the activity of *Corvus corone cornix* along the day. The x-axis refers to the time of day. The y-axis is the date. The numbers in parentheses for dates represent the total number of events detected during the day. The orange line separates lockdown from no lockdown periods.

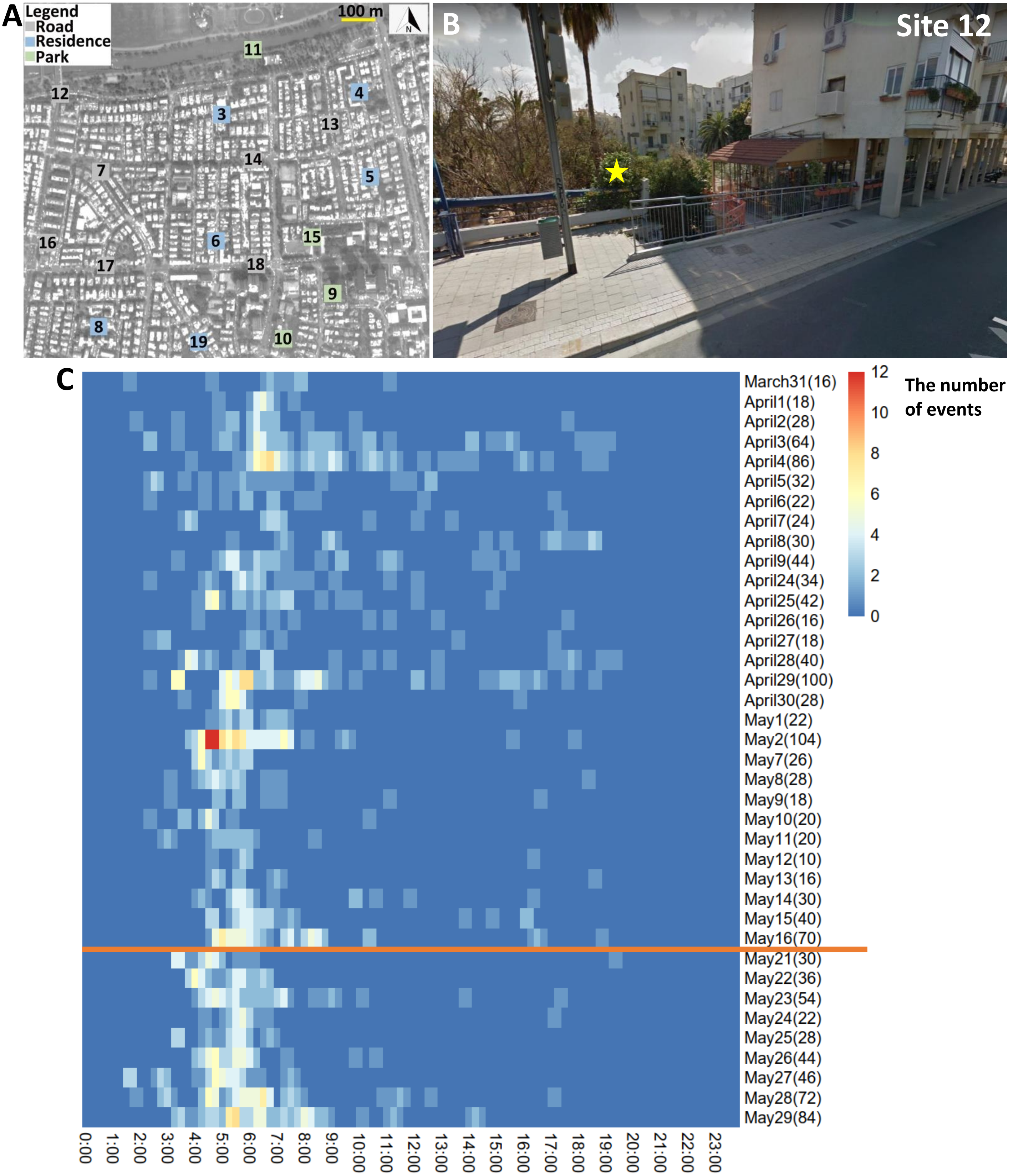


**Figure S10.** (A) Study area and (B) recording site 12. The yellow star refers to the audiomoth's location. (C) Heatmap indicating the activity of *Corvus corone cornix* along the day. The x-axis refers to the time of day. The y-axis is the date. The numbers in parentheses for dates represent the total number of events detected during the day. The orange line separates lockdown from no lockdown periods.

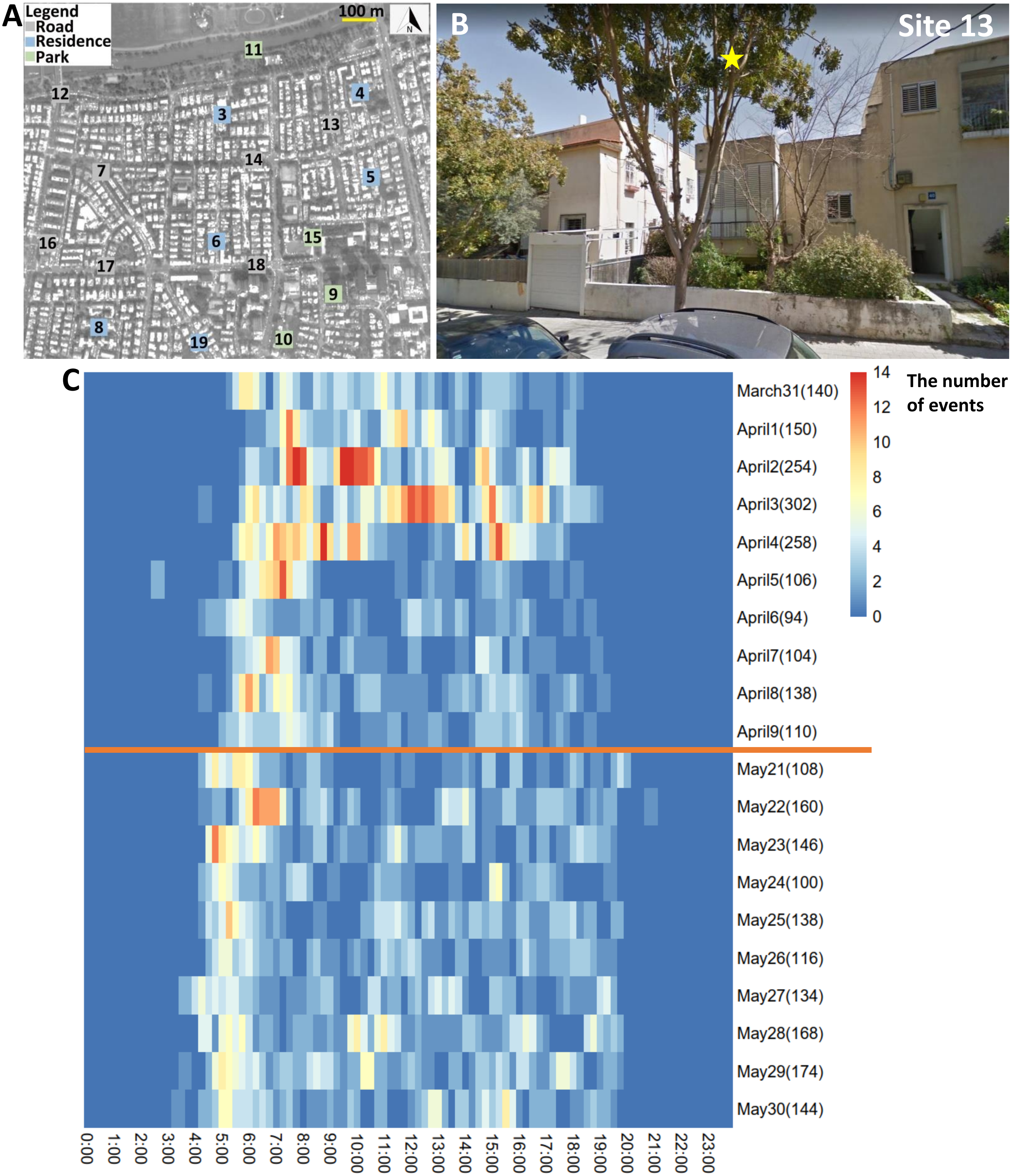


**Figure S11.** (A) Study area and (B) recording site 13. The yellow star refers to the audiomoth's location. (C) Heatmap indicating the activity of *Corvus corone cornix* along the day. The x-axis refers to the time of day. The y-axis is the date. The numbers in parentheses for dates represent the total number of events detected during the day. The orange line separates lockdown from no lockdown periods.

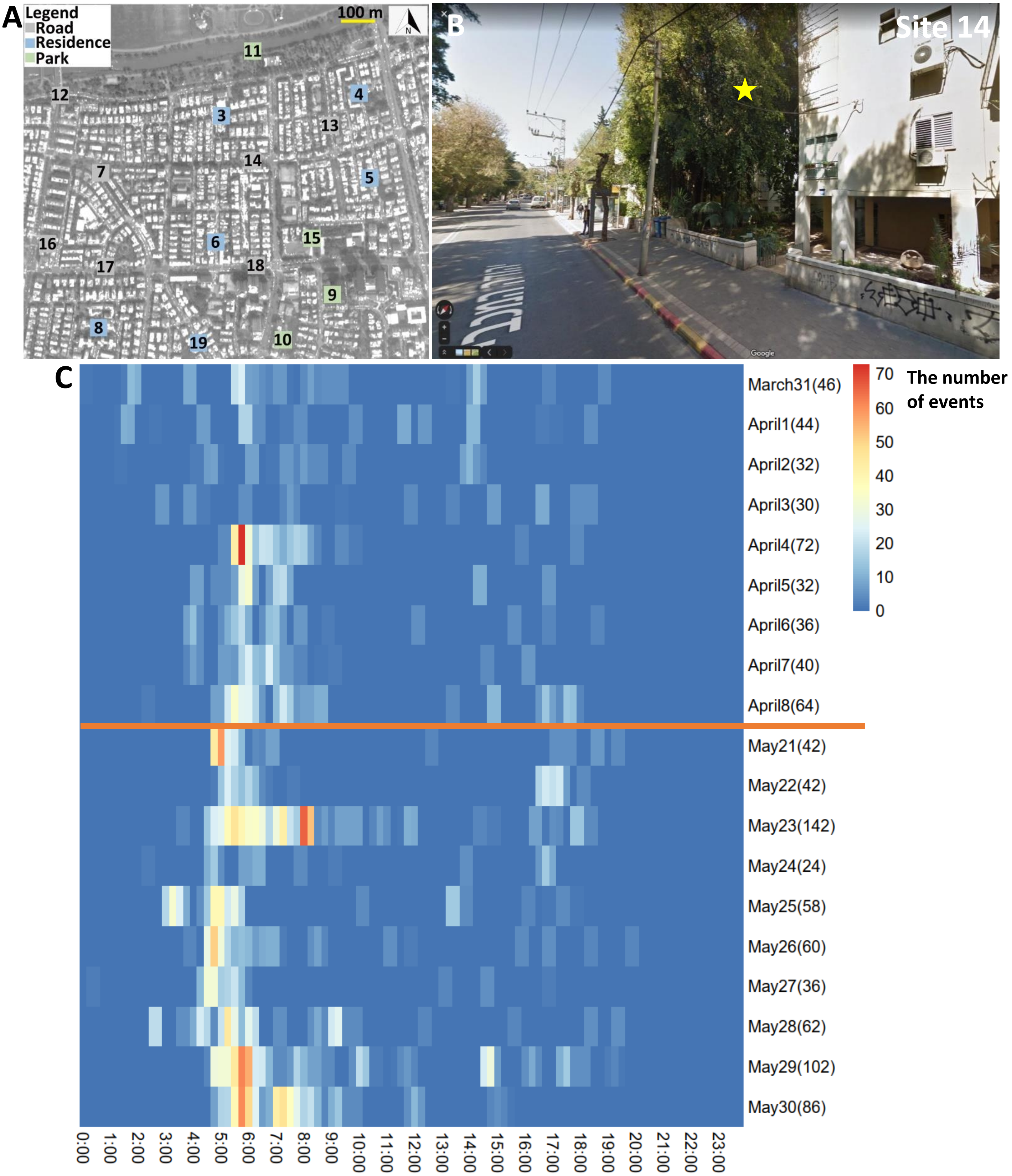


**Figure S12.** (A) Study area and (B) recording site 14. The yellow star refers to the audiomoth's location. (C) Heatmap indicating the activity of *Corvus corone cornix* along the day. The x-axis refers to the time of day. The y-axis is the date. The numbers in parentheses for dates represent the total number of events detected during the day. The orange line separates lockdown from no lockdown periods.

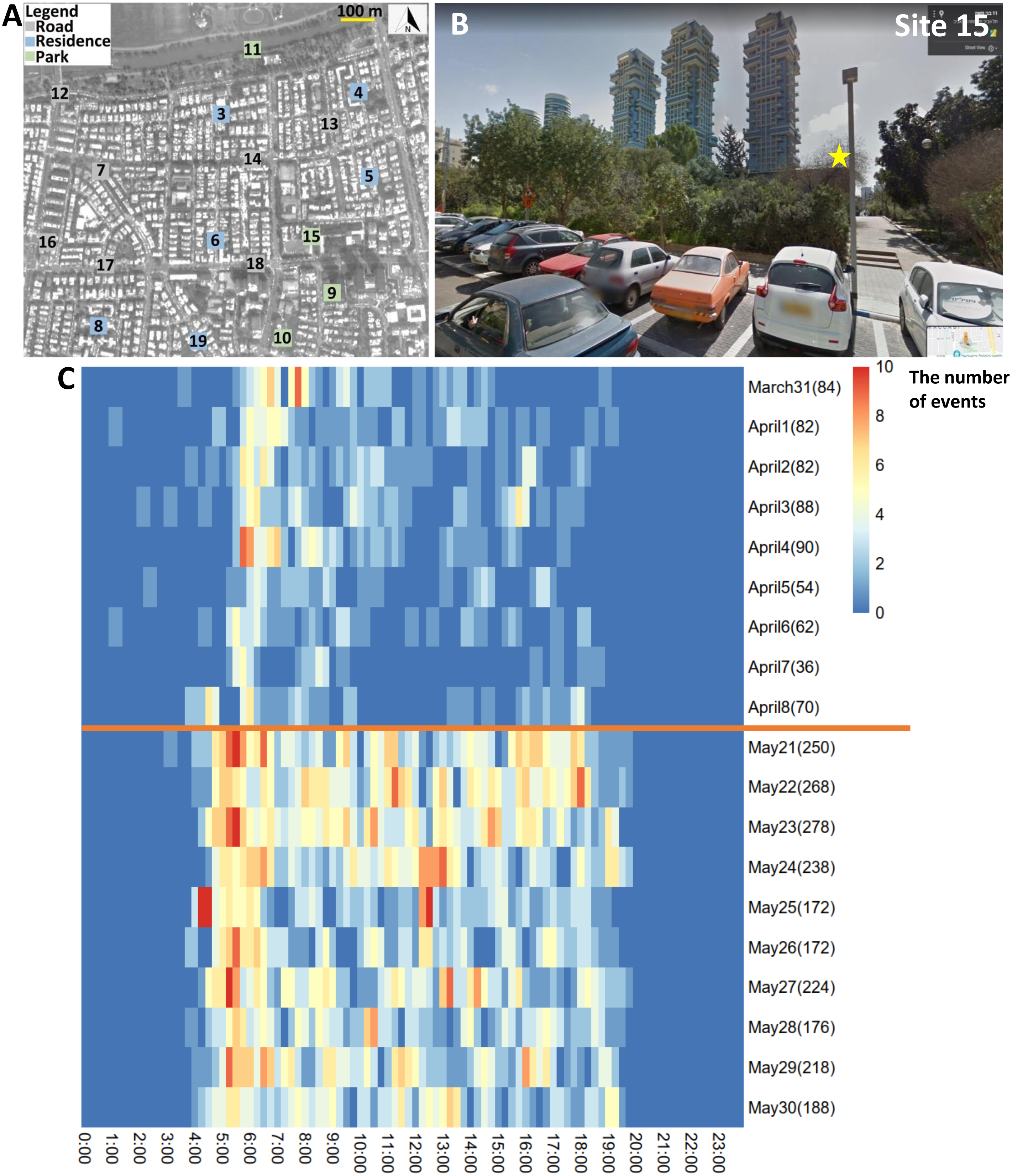


**Figure S13.** (A) Study area and (B) recording site 15. The yellow star refers to the audiomoth's location. (C) Heatmap indicating the activity of *Corvus corone cornix* along the day. The x-axis refers to the time of day. The y-axis is the date. The numbers in parentheses for dates represent the total number of events detected during the day. The orange line separates lockdown from no lockdown periods.

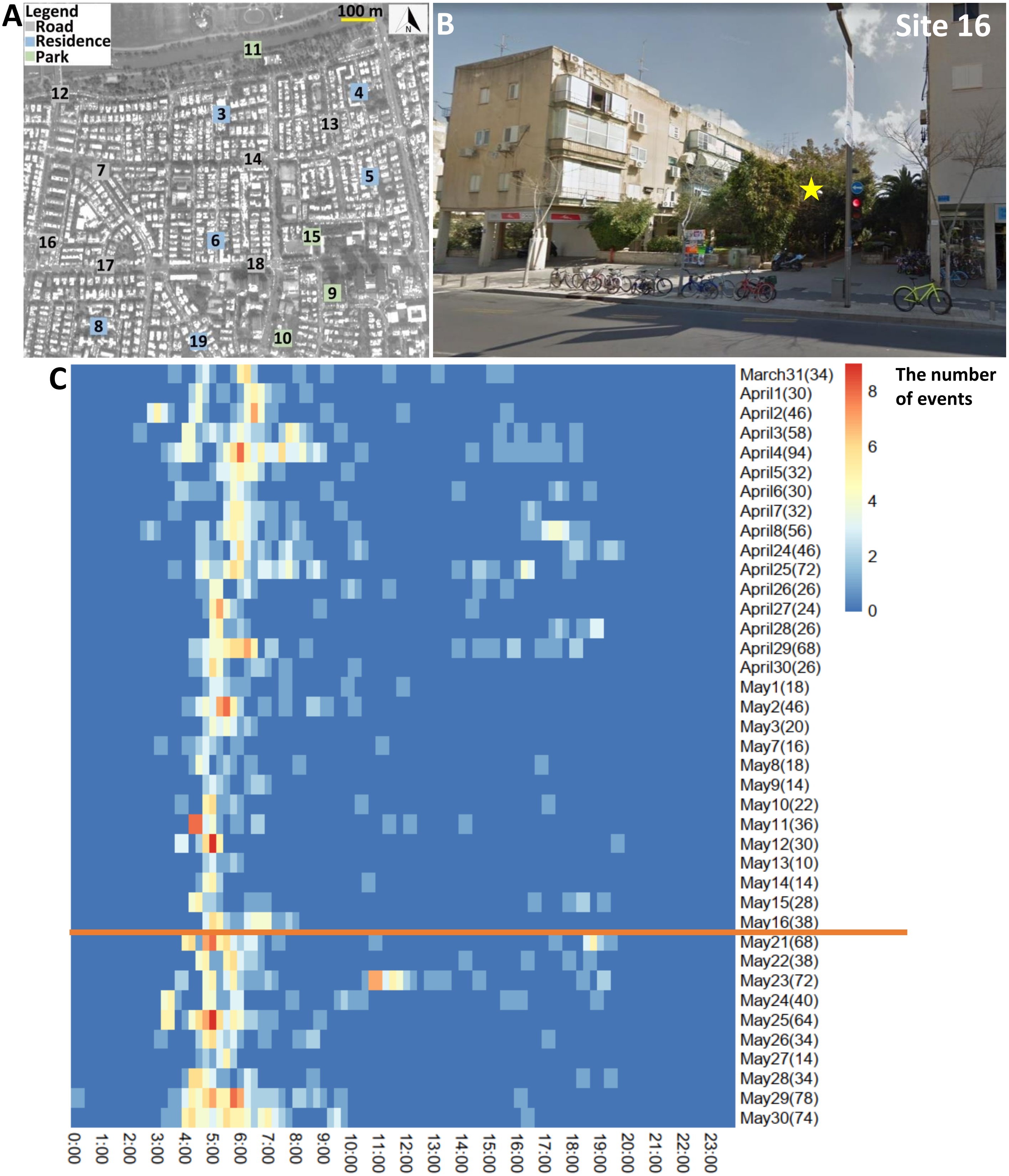


**Figure S14.** (A) Study area and (B) recording site 16. The yellow star refers to the audiomoth's location. (C) Heatmap indicating the activity of *Corvus corone cornix* along the day. The x-axis refers to the time of day. The y-axis is the date. The numbers in parentheses for dates represent the total number of events detected during the day. The orange line separates lockdown from no lockdown periods.

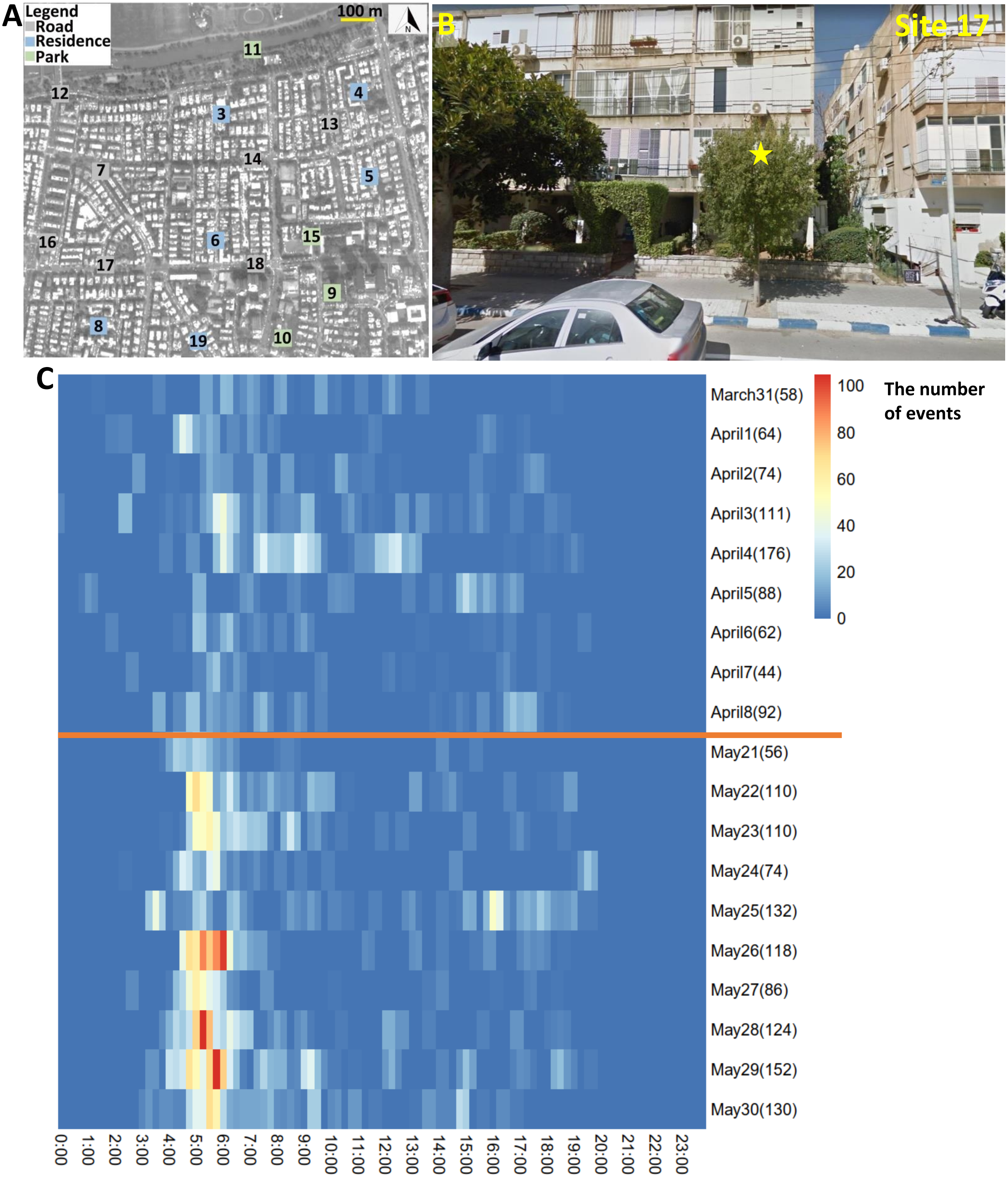


**Figure S15.** (A) Study area and (B) recording site 17. The yellow star refers to the audiomoth's location. (C) Heatmap indicating the activity of *Corvus corone cornix* along the day. The x-axis refers to the time of day. The y-axis is the date. The numbers in parentheses for dates represent the total number of events detected during the day. The orange line separates lockdown from no lockdown periods.

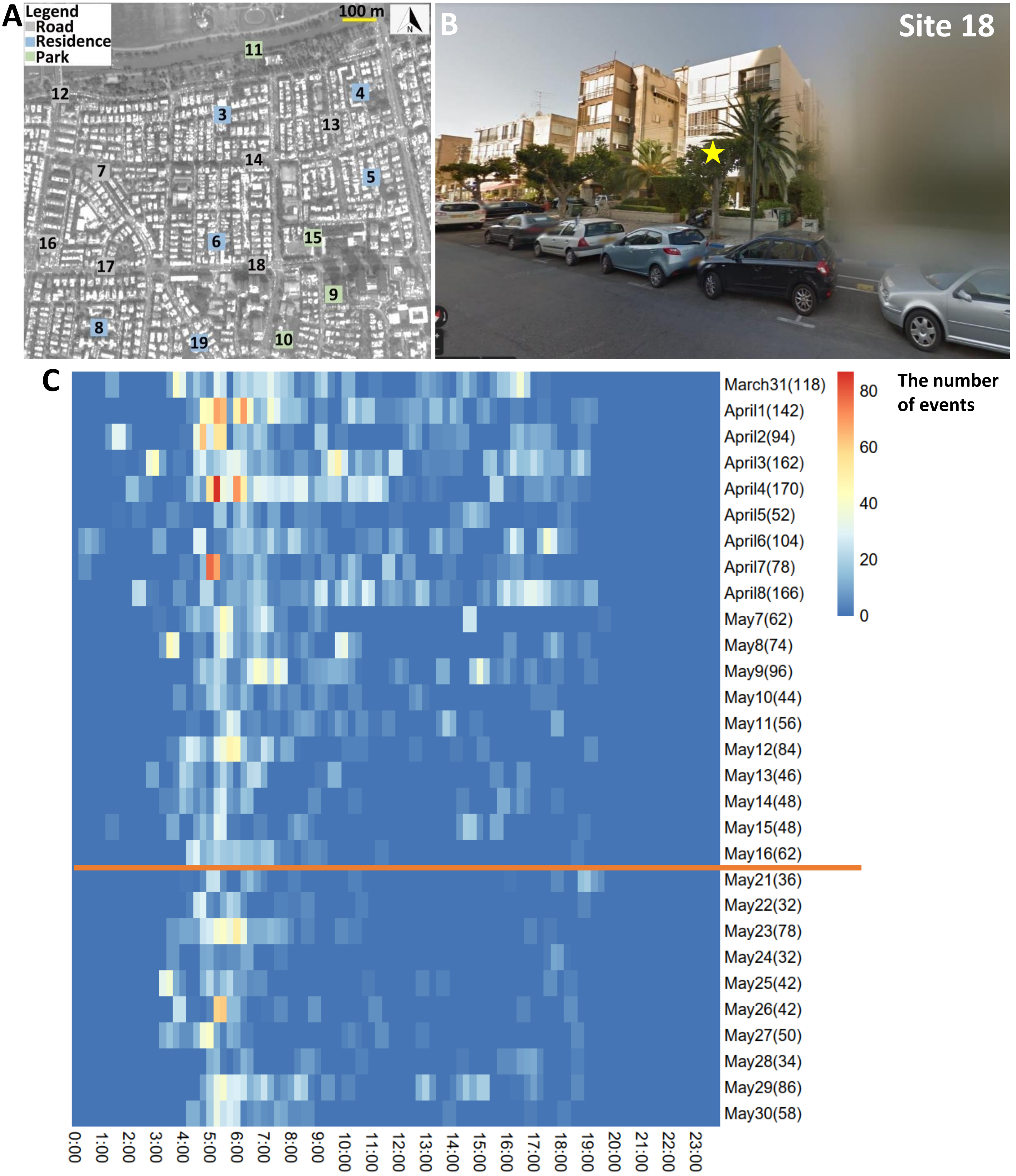


**Figure S16.** (A) Study area and (B) recording site 18. The yellow star refers to the audiomoth's location. (C) Heatmap indicating the activity of *Corvus corone cornix* along the day. The x-axis refers to the time of day. The y-axis is the date. The numbers in parentheses for dates represent the total number of events detected during the day. The orange line separates lockdown from no lockdown periods.

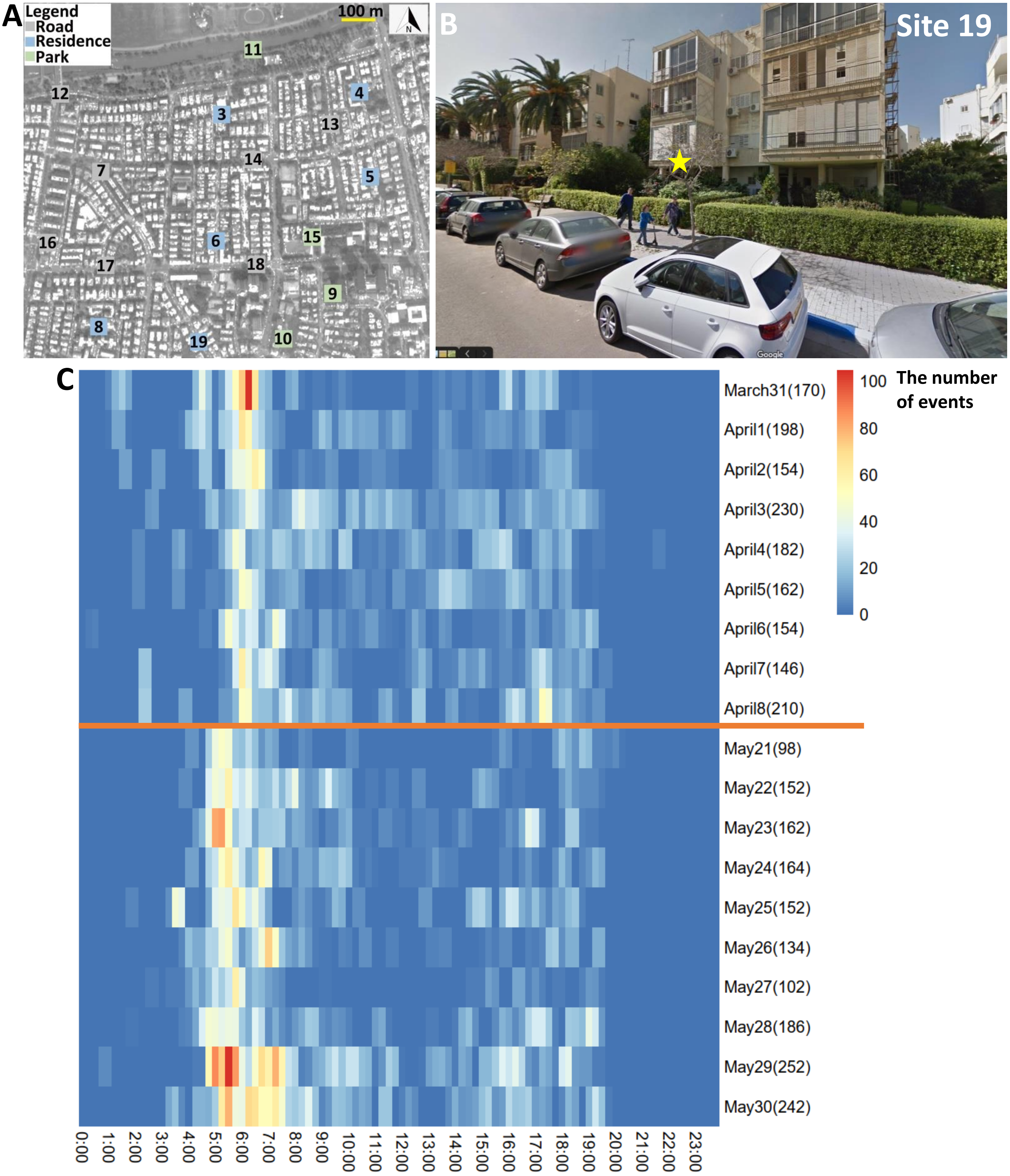


**Figure S17.** (A) Study area and (B) recording site 19. The yellow star refers to the audiomoth's location. (C) Heatmap indicating the activity of *Corvus corone cornix* along the day. The x-axis refers to the time of day. The y-axis is the date. The numbers in parentheses for dates represent the total number of events detected during the day. The orange line separates lockdown from no lockdown periods.
