## Supplemental Figure S1-S7, Table S6-S19, and will be used for the link to the file on the preprint site. for "Species and habitat specific changes in bird activity in an urban environment during Covid 19 lockdown"

**Table S6** Results from generalized linear mixed models (GLMM) comparing the differences in human activity between no lockdown period and lockdown period for all sites and for each site.

| Dependent variable | Site category | Estimate | *P* | Percent |
| --- | --- | --- | --- | --- |
| Human_activity | All sites | -0.444 | **< 0.0001** | 35.85 |
|  | Road | 0.082 | 0.632 | 8.55 |
|  | Residence | 0.398 | **< 0.0001** | 48.88 |
|  | Park | -0.375 | **0.002** | 31.27 |

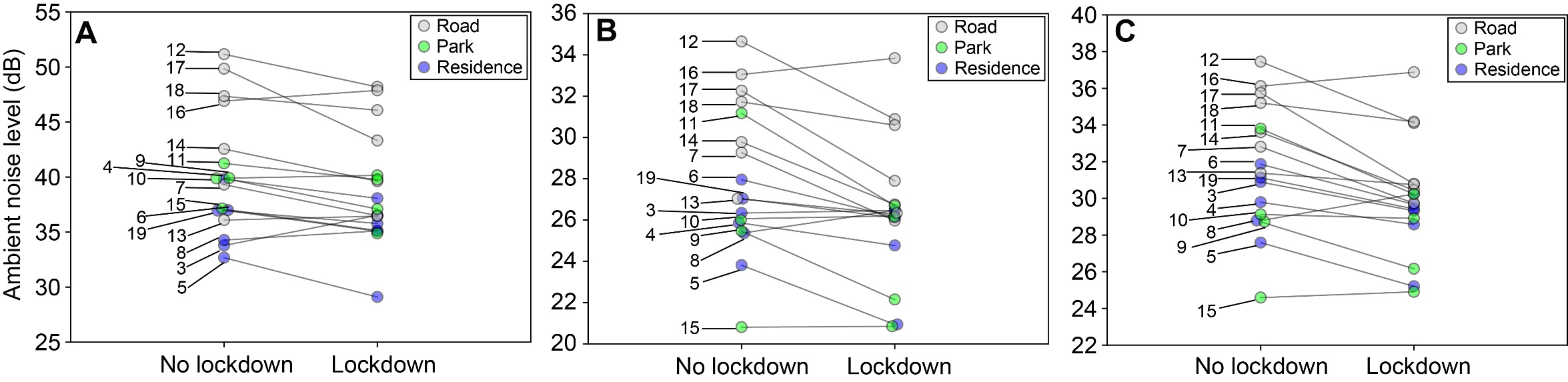

**Figure S1** The ambient noise level **(**dB**)** between no lockdown period and lockdown period.

**Table S7** Akaike’s information criterion (AICc) model comparison results for activity and activity variability in all species and each bird species.

| Species | Model description | *df* | logLik | AICc | ∆AICc | *wi* |
| --- | --- | --- | --- | --- | --- | --- |
| **All species** | **Activity (Number of events/day)** ~ |  |  |  |  |  |
|  | Bird, Lock, Noise, Site, Human, Temperature, Bird*Lock, Count*Lock, Lock*Noise, Lock*Site, Lock*Human | 16 | -29807.60 | 59647.53 | 0.00 | 0.93 |
|  | Bird, Lock, Noise, Site, Human, Temperature, Bird*Lock, Count*Lock, Lock*Site, Lock*Human | 15 | -29811.28 | 59652.86 | 5.33 | 0.07 |
| **All species** | **Activity Variability (CV of the number of events/day) ~** |  |  |  |  |  |
|  | Bird, Lock, Noise, Site, Human, Temperature, Bird*Lock, Count*Lock, Lock*Noise | 14 | 235.75 | -443.21 | 0.00 | 0.22 |
|  | Bird, Lock, Noise, Site, Human, Temperature, Bird*Lock, Count*Lock, Lock*Noise, Lock*Human | 15 | 236.70 | -443.06 | 0.14 | 0.20 |
|  | Bird, Lock, Noise, Site, Human, Temperature, Bird*Lock, Lock*Noise | 13 | 234.60 | -442.96 | 0.25 | 0.19 |
|  | Bird, Lock, Noise, Site, Human, Temperature, Bird*Lock, Lock*Noise, Lock*Human | 14 | 235.53 | -442.78 | 0.43 | 0.17 |
|  | Bird, Lock, Noise, Site, Human, Temperature, Bird*Lock, Lock*Noise, Lock*Site | 15 | 235.48 | -440.62 | 2.58 | 0.06 |
|  | Bird, Lock, Noise, Site, Human, Temperature, Bird*Lock, Count*Lock, Lock*Noise, Lock*Site | 16 | 236.38 | -440.40 | 2.81 | 0.05 |
|  | Bird, Lock, Noise, Site, Temperature, Bird*Lock, Count*Lock, Lock*Noise | 13 | 233.07 | -439.89 | 3.32 | 0.04 |
|  | Bird, Lock, Noise, Site, Human, Temperature, Bird*Lock, Lock*Noise, Lock*Site, Lock*Human | 16 | 235.97 | -439.56 | 3.65 | 0.03 |
|  | Bird, Lock, Noise, Site, Human, Temperature, Bird*Lock, Count*Lock, Lock*Noise, Lock*Site, Lock*Human | 17 | 236.93 | -439.43 | 3.78 | 0.03 |
| **Hooded crow** | **Activity (Number of events/day)** ~ |  |  |  |  |  |
|  | Lock, Noise, Site, Human, Count*Lock, Lock*Site, Lock*Human | 12 | -5327.10 | 10678.79 | 0.00 | 0.48 |
|  | Lock, Noise, Site, Human, Count*Lock, Lock*Noise, Lock*Site, Lock*Human | 13 | -5326.86 | 10680.41 | 1.61 | 0.22 |
|  | Lock, Noise, Site, Human, Temperature, Count*Lock, Lock*Site, Lock*Human | 13 | -5326.88 | 10680.46 | 1.67 | 0.21 |
|  | Lock, Noise, Site, Human, Temperature, Count*Lock, Lock*Noise, Lock*Site, Lock*Human | 14 | -5326.67 | 10682.15 | 3.36 | 0.09 |
| **Hooded crow** | **Activity Variability (CV of the number of events/day) ~** |  |  |  |  |  |
|  | Lock, Noise, Site, Human, Lock*Noise, Lock*Human | 11 | 247.29 | -472.08 | 0.00 | 0.28 |
|  | Lock, Noise, Site, Human, Temperature, Lock*Noise, Lock*Human | 12 | 248.02 | -471.44 | 0.64 | 0.21 |
|  | Lock, Noise, Site, Human, Count*Lock, Lock*Noise, Lock*Human | 12 | 247.94 | -471.29 | 0.79 | 0.19 |
|  | Lock, Noise, Site, Human, Temperature, Count*Lock, Lock*Noise, Lock*Human | 13 | 248.26 | -469.83 | 2.25 | 0.09 |
|  | Lock, Noise, Site, Human, Lock*Noise, Lock*Site, Lock*Human | 13 | 247.99 | -469.29 | 2.79 | 0.07 |
|  | Lock, Noise, Site, Human, Count*Lock, Lock*Noise, Lock*Site, Lock*Human | 14 | 248.92 | -469.05 | 3.03 | 0.06 |
|  | Lock, Noise, Site, Human, Temperature, Lock*Noise, Lock*Site, Lock*Human | 14 | 248.69 | -468.58 | 3.50 | 0.05 |
|  | Lock, Noise, Human, Lock*Noise, Lock*Human | 9 | 243.29 | -468.23 | 3.85 | 0.04 |
| **Rose-ringed**  **parakeet** | **Activity (Number of events/day)** ~ |  |  |  |  |  |
|  | Lock, Noise, Site, Human, Temperature, Lock*Noise, Lock*Site, Lock*Human | 13 | -3631.41 | 7289.51 | 0.00 | 0.60 |
|  | Lock, Noise, Site, Human, Temperature, Count*Lock, Lock*Noise, Lock*Site, Lock*Human | 14 | -3630.81 | 7290.43 | 0.91 | 0.38 |
|  | Lock, Noise, Site, Human, Temperature, Count*Lock, Lock*Site, Lock*Human | 13 | -3635.75 | 7298.19 | 8.67 | 0.01 |
|  | Lock, Noise, Site, Human, Temperature, Lock*Site, Lock*Human | 12 | -3636.94 | 7298.47 | 8.96 | 0.01 |
| **Rose-ringed**  **parakeet** | **Activity Variability (CV of the number of events/day) ~** |  |  |  |  |  |
|  | Lock, Noise, Site, Human, Temperature, Count*Lock, Lock*Site, Lock*Human | 14 | 462.79 | -896.76 | 0.00 | 0.58 |
|  | Lock, Noise, Site, Human, Temperature, Count*Lock, Lock*Noise, Lock*Site, Lock*Human | 15 | 463.14 | -895.35 | 1.41 | 0.29 |
|  | Lock, Noise, Site, Human, Count*Lock, Lock*Site, Lock*Human | 13 | 460.23 | -893.76 | 3.00 | 0.13 |
| **Graceful prinia** | **Activity (Number of events/day)** ~ |  |  |  |  |  |
|  | Lock, Noise, Site, Human, Temperature, Count*Lock, Lock*Noise, Lock*Site, Lock*Human | 14 | -8002.31 | 16033.42 | 0.00 | 0.99 |
|  | Lock, Noise, Site, Human, Temperature, Count*Lock, Lock*Noise, Lock*Site | 13 | -8008.61 | 16043.92 | 10.5 | 0.01 |
| **Graceful prinia** | **Activity Variability (CV of the number of events/day) ~** |  |  |  |  |  |
|  | Lock, Noise, Site, Human, Temperature, Count*Lock, Lock*Site | 12 | -227.65 | 480.08 | 0.00 | 0.11 |
|  | Lock, Noise, Site, Temperature | 9 | -230.97 | 480.40 | 0.32 | 0.09 |
|  | Lock, Noise, Site, Temperature, Lock*Site | 11 | -228.89 | 480.44 | 0.36 | 0.09 |
|  | Lock, Noise, Site, Temperature, Count*Lock | 10 | -230.01 | 480.57 | 0.49 | 0.09 |
|  | Lock, Site, Temperature, Count*Lock, Lock*Site | 11 | -229.3 | 481.26 | 1.18 | 0.06 |
|  | Noise, Site, Temperature | 8 | -232.53 | 481.42 | 1.34 | 0.06 |
|  | Lock, Noise, Site, Temperature, Count*Lock, Lock*Noise, Lock*Site | 13 | -227.44 | 481.81 | 1.73 | 0.05 |
|  | Lock, Noise, Site, Human, Temperature, Count*Lock, Lock*Site | 13 | -227.45 | 481.83 | 1.75 | 0.05 |
|  | Lock, Site, Temperature, Count*Lock | 9 | -231.73 | 481.92 | 1.84 | 0.04 |
|  | Lock, Noise, Site, Human, Temperature, Lock*Site | 12 | -228.76 | 482.32 | 2.24 | 0.04 |
|  | Lock, Noise, Site, Temperature, Lock*Noise, Lock*Site | 12 | -228.82 | 482.43 | 2.36 | 0.03 |
|  | Lock, Site, Temperature, Lock*Site | 10 | -230.96 | 482.48 | 2.40 | 0.03 |
|  | Lock, Noise, Site, Human, Temperature | 10 | -230.97 | 482.49 | 2.41 | 0.03 |
|  | Lock, Noise, Site, Temperature, Lock*Noise | 10 | -230.97 | 482.49 | 2.42 | 0.03 |
|  | Lock, Site, Temperature | 8 | -233.12 | 482.59 | 2.52 | 0.03 |
|  | Lock, Noise, Site, Human, Temperature, Count*Lock | 11 | -229.99 | 482.64 | 2.56 | 0.03 |
|  | Lock, Noise, Site, Temperature, Count*Lock, Lock*Noise | 11 | -229.99 | 482.65 | 2.57 | 0.03 |
|  | Lock, Site, Human, Temperature, Count*Lock, Lock*Site | 12 | -229.02 | 482.83 | 2.75 | 0.03 |
|  | Lock, Noise, Site, Human, Temperature, Count*Lock, Lock*Noise, Lock*Site | 14 | -227.20 | 483.46 | 3.38 | 0.02 |
|  | Noise, Site, Human, Temperature | 9 | -232.51 | 483.48 | 3.40 | 0.02 |
|  | Lock, Noise, Site, Human, Temperature, Count*Lock, Lock*Site, Lock*Human | 14 | -227.38 | 483.82 | 3.74 | 0.02 |
|  | Lock, Site, Human, Temperature, Count*Lock | 10 | -231.69 | 483.93 | 3.85 | 0.02 |

Bird: Bird species. Lock: Lockdown status. Site: Site category. Human: Human activity. Count: Count down. AICc: Akaike’s information criterion corrected for small sample size. ΔAICc: the difference between the alternative model and best-fitting model. Models are ranked based on the AICc values from the best to the worst model. *: Interaction effect.

**Table S8** Results from post hoc tests comparing the differences in activity and activity variability between no lockdown period and lockdown period in all species for each site. Estimates were calculated in % per day for the following units: Temp – per degree, Noise – per dB, Human activity – per 1 talking event, Lockdown related parameter – per existence of the lockdown (yes/no).

| Species | Dependent variable | Site category | Estimate | *P* | Percent |
| --- | --- | --- | --- | --- | --- |
| All species | **Activity**-  Number of events/day | Road | -0.136 | 0.230 | 12.716 |
|  |  | Residence | 0.029 | 0.800 | 2.942 |
|  |  | Park | -0.043 | 0.707 | 4.209 |
|  | **Activity variability**-  CV of the number of events/day | Road | -0.037 | 0.260 | - |
|  |  | Residence | -0.016 | 0.663 | - |
|  |  | Park | -0.001 | 0.988 | - |
| Hooded crow | **Activity**-  Number of events/day | Road | 0.032 | 0.828 | 3.252 |
|  |  | Residence | 0.328 | **0.025** | 38.819 |
|  |  | Park | 0.258 | 0.076 | 29.434 |
|  | **Activity variability**-  CV of the number of events/day | Road | -0.046 | 0.217 | - |
|  |  | Residence | -0.078 | 0.080 | - |
|  |  | Park | -0.034 | 0.461 | - |
| Rose-ringed parakeet | **Activity**-  Number of events/day | Road | 0.066 | 0.664 | 6.823 |
|  |  | Residence | 0.242 | 0.114 | 27.379 |
|  |  | Park | 0.643 | **< 0.0001** | 90.218 |
|  | **Activity variability**-  CV of the number of events/day | Road | -0.029 | 0.332 | - |
|  |  | Residence | -0.070 | **0.038** | - |
|  |  | Park | -0.128 | **0.0003** | - |
| Graceful prinia | **Activity-**  Number of events/day | Road | -0.440 | 0.511 | 2.858 |
|  |  | Residence | -1.580 | **0.018** | 6.761 |
|  |  | Park | -1.640 | **0.014** | 12.015 |
|  | **Activity variability**-  CV of the number of events/day | Road | -0.038 | 0.781 | - |
|  |  | Residence | 0.142 | 0.285 | - |
|  |  | Park | 0.303 | **0.030** | - |

CV: coefficient of variance

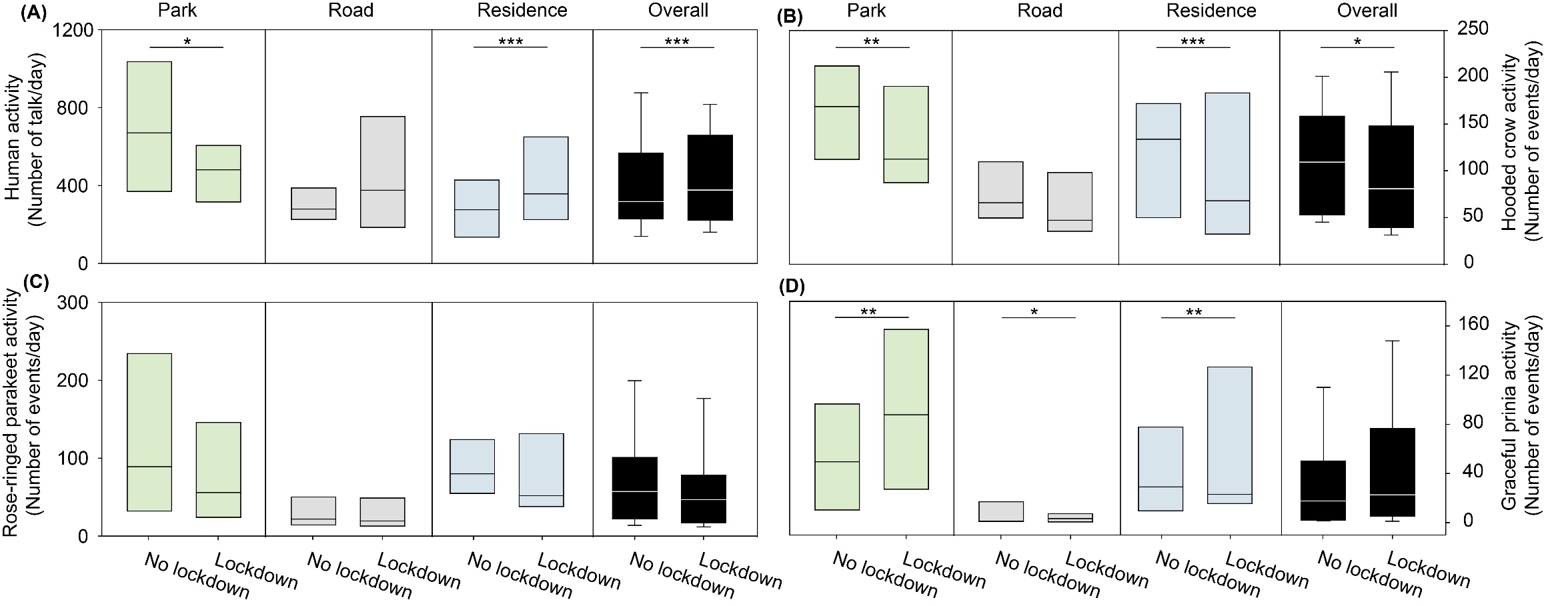

**Figure S2. Activity of birds**. Boxplots show the activity of (A) Human activity, (B) Hooded crow*,* (C) Rose-ringed parakeet, and (D) Graceful prinia for each site category during no lockdown and during lockdown. Green box plot: park; Grey box plot: road; Blue box plot: residence; Black box plot: overall. Note that human activity was assessed based on human speech so the increase observed during lockdowns in roads represents pedestrian and not car activity. Box plot lower and upper box boundaries show the 25th and 75th percentiles, respectively, with the median inside. The lower and upper error lines depict the 10th and 90th percentiles, respectively. Outliers of the data are shown as black dots. **P* < 0.05, ***P* < 0.01, ****P* < 0.001.

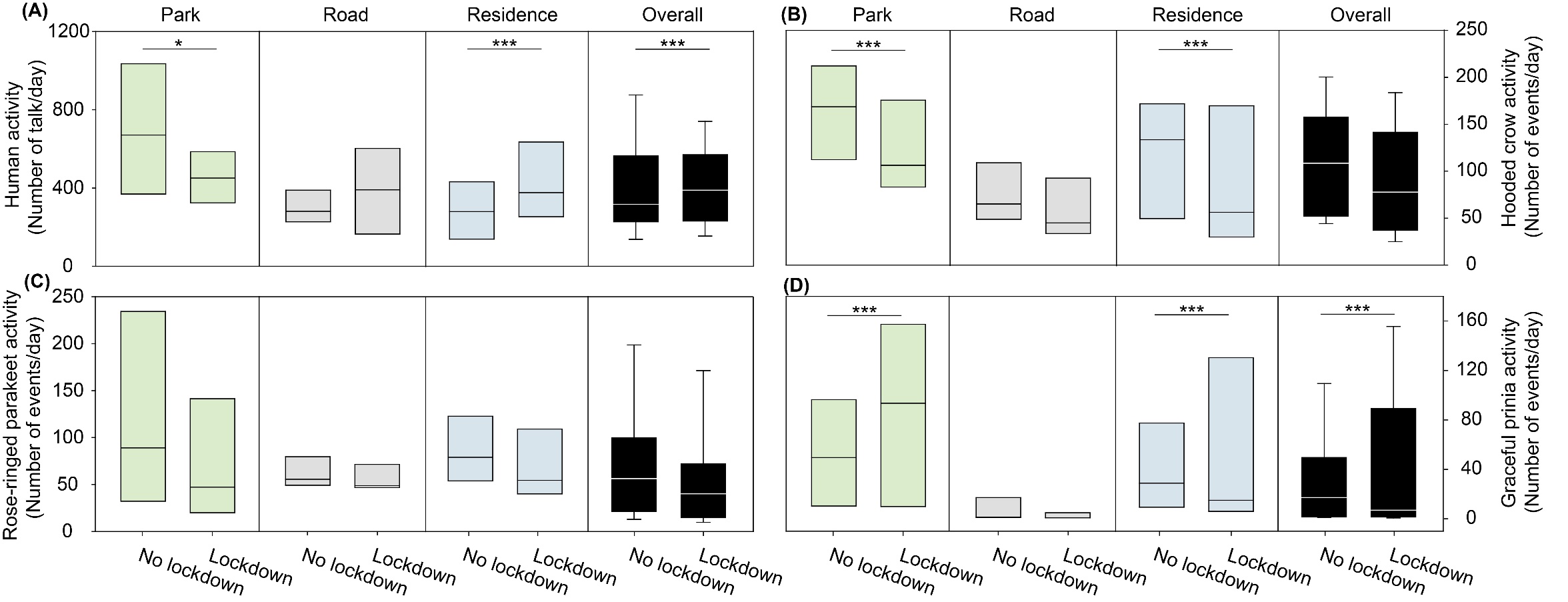

**Figure S3. Activity of birds**. Boxplots show the activity of (A) Human activity, (B) Hooded crow*,* (C) Rose-ringed parakeet, and (D) Graceful prinia for each site category during no lockdown and during lockdown. Green box plot: park; Grey box plot: road; Blue box plot: residence; Black box plot: overall. Note that human activity was assessed based on human speech so the increase observed during lockdowns in roads represents pedestrian and not car activity. Box plot lower and upper box boundaries show the 25th and 75th percentiles, respectively, with the median inside. The lower and upper error lines depict the 10th and 90th percentiles, respectively. Outliers of the data are shown as black dots. **P* < 0.05, ***P* < 0.01, ****P* < 0.001.

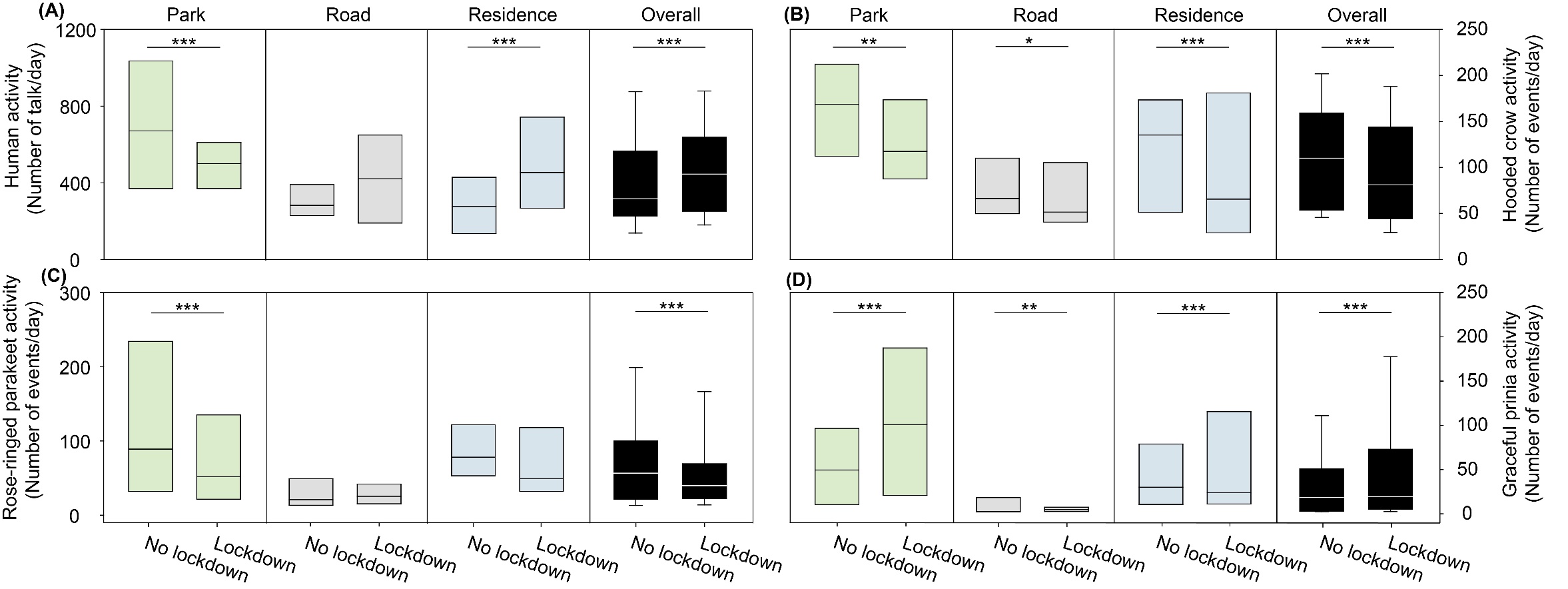

**Figure S4. Activity of birds**. Boxplots show the activity of (A) Human activity, (B) Hooded crow*,* (C) Rose-ringed parakeet, and (D) Graceful prinia for each site category during no lockdown and during lockdown. Green box plot: park; Grey box plot: road; Blue box plot: residence; Black box plot: overall. Note that human activity was assessed based on human speech so the increase observed during lockdowns in roads represents pedestrian and not car activity. Box plot lower and upper box boundaries show the 25th and 75th percentiles, respectively, with the median inside. The lower and upper error lines depict the 10th and 90th percentiles, respectively. Outliers of the data are shown as black dots. **P* < 0.05, ***P* < 0.01, ****P* < 0.001.

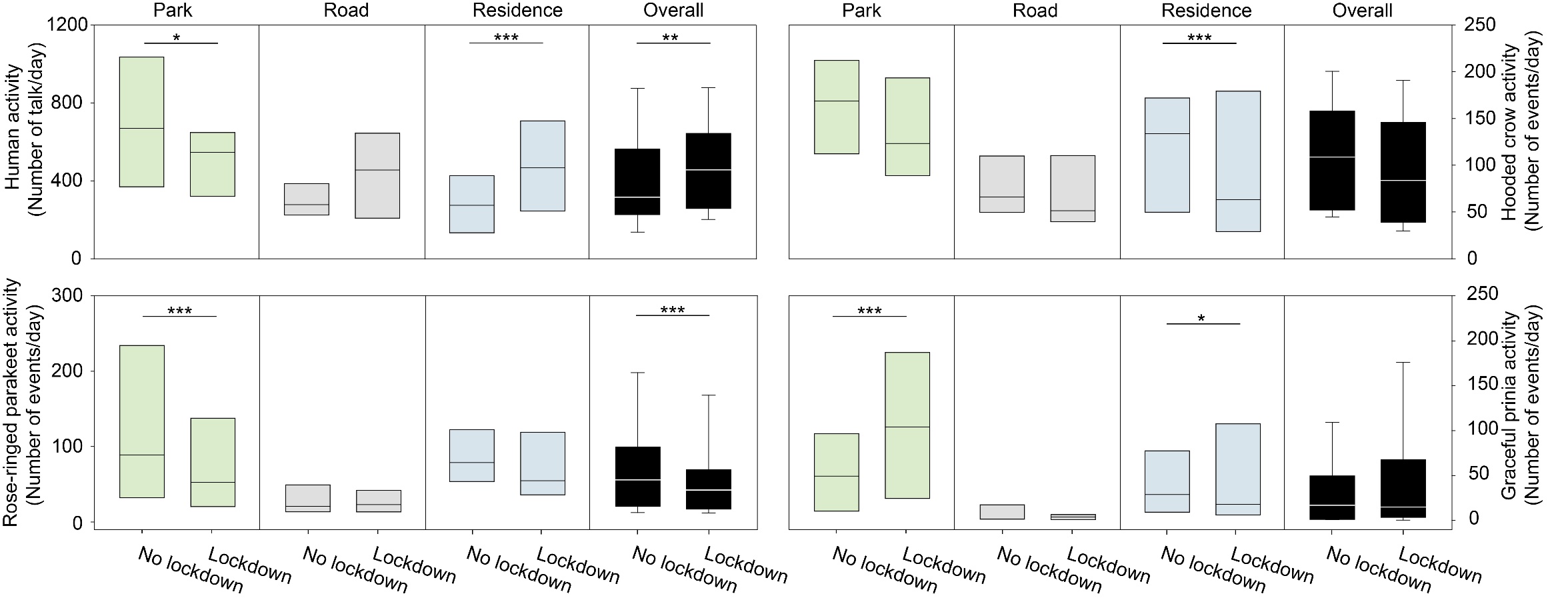

**Figure S5. Activity of birds**. Boxplots show the activity of (A) Human activity, (B) Hooded crow*,* (C) Rose-ringed parakeet, and (D) Graceful prinia for each site category during no lockdown and during lockdown. Green box plot: park; Grey box plot: road; Blue box plot: residence; Black box plot: overall. Note that human activity was assessed based on human speech so the increase observed during lockdowns in roads represents pedestrian and not car activity. Box plot lower and upper box boundaries show the 25th and 75th percentiles, respectively, with the median inside. The lower and upper error lines depict the 10th and 90th percentiles, respectively. Outliers of the data are shown as black dots. **P* < 0.05, ***P* < 0.01, ****P* < 0.001.

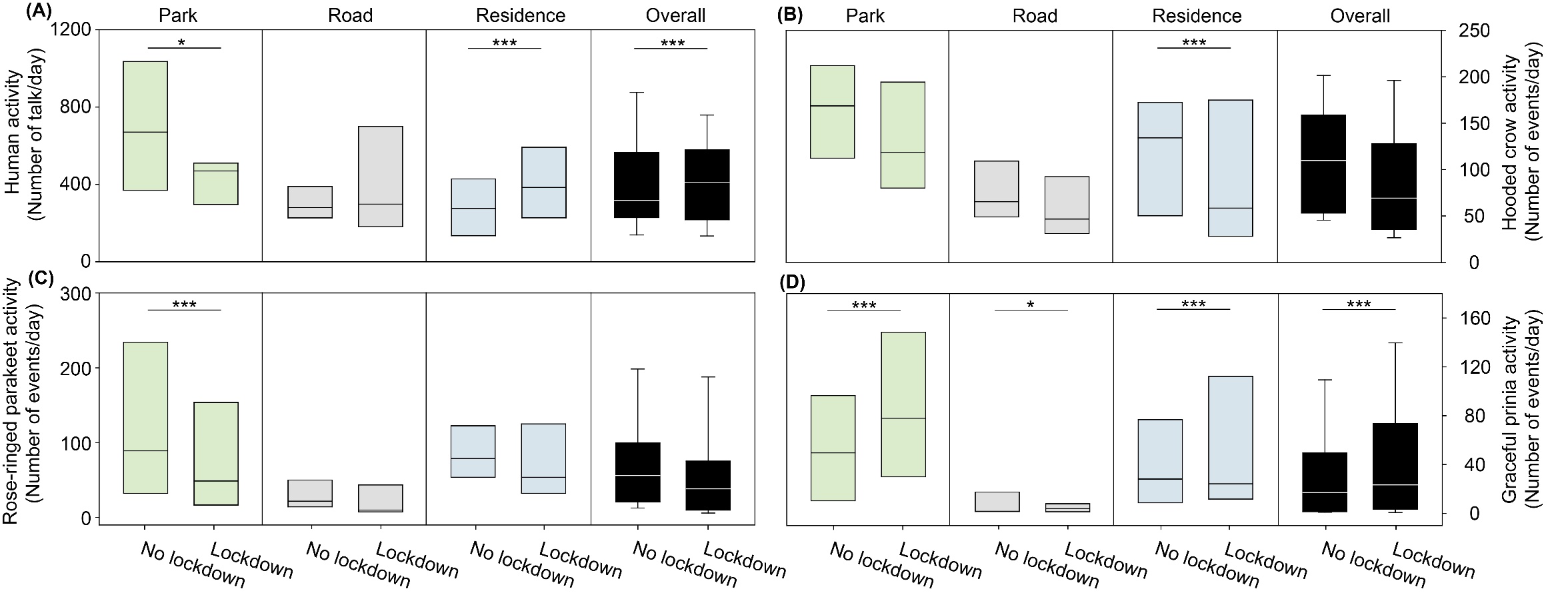

**Figure S6. Activity of birds**. Boxplots show the activity of (A) Human activity, (B) Hooded crow*,* (C) Rose-ringed parakeet, and (D) Graceful prinia for each site category during no lockdown and during lockdown. Green box plot: park; Grey box plot: road; Blue box plot: residence; Black box plot: overall. Note that human activity was assessed based on human speech so the increase observed during lockdowns in roads represents pedestrian and not car activity. Box plot lower and upper box boundaries show the 25th and 75th percentiles, respectively, with the median inside. The lower and upper error lines depict the 10th and 90th percentiles, respectively. Outliers of the data are shown as black dots. **P* < 0.05, ***P* < 0.01, ****P* < 0.001.

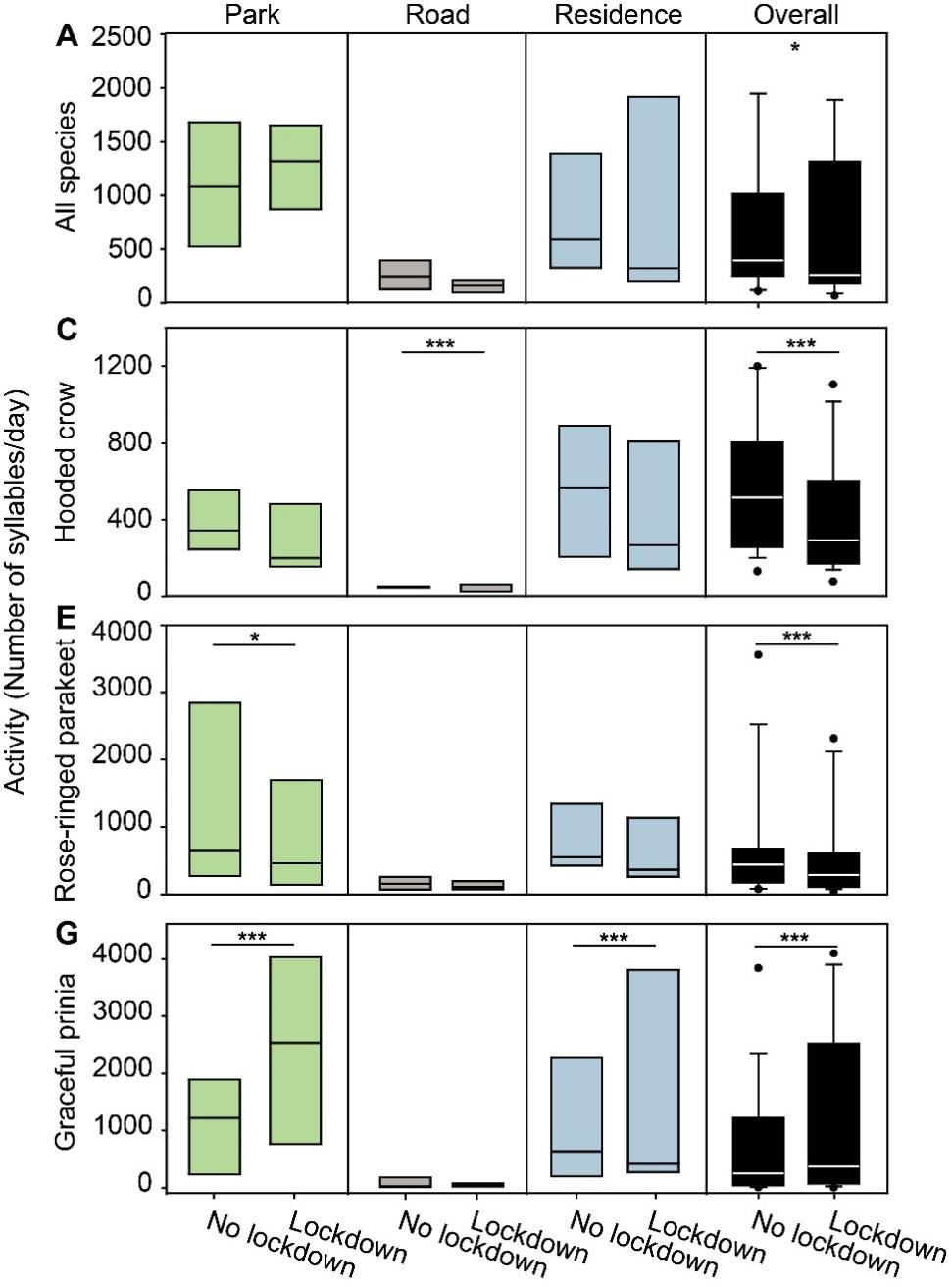

**Figure S7. Activity of birds**. Boxplots show the activity of (A) all species, (B) Hooded crow, (C) Rose-ringed parakeet, and (D) Graceful prinia for each site category during no lockdown and during lockdown. Green box plot: park; Grey box plot: road; Blue box plot: residence; Black box plot: overall. Box plot lower and upper box boundaries show the 25th and 75th percentiles, respectively, with the median inside. The lower and upper error lines depict the 10th and 90th percentiles, respectively. Outliers of the data are shown as black dots. Asterisks indicate significant differences between no lockdown and lockdown periods. **P* < 0.05, ***P* < 0.01, ****P* < 0.001. Activity is defined as the daily number of syllables of bird vocalizations per day. Different sampling sites represented by different symbols.

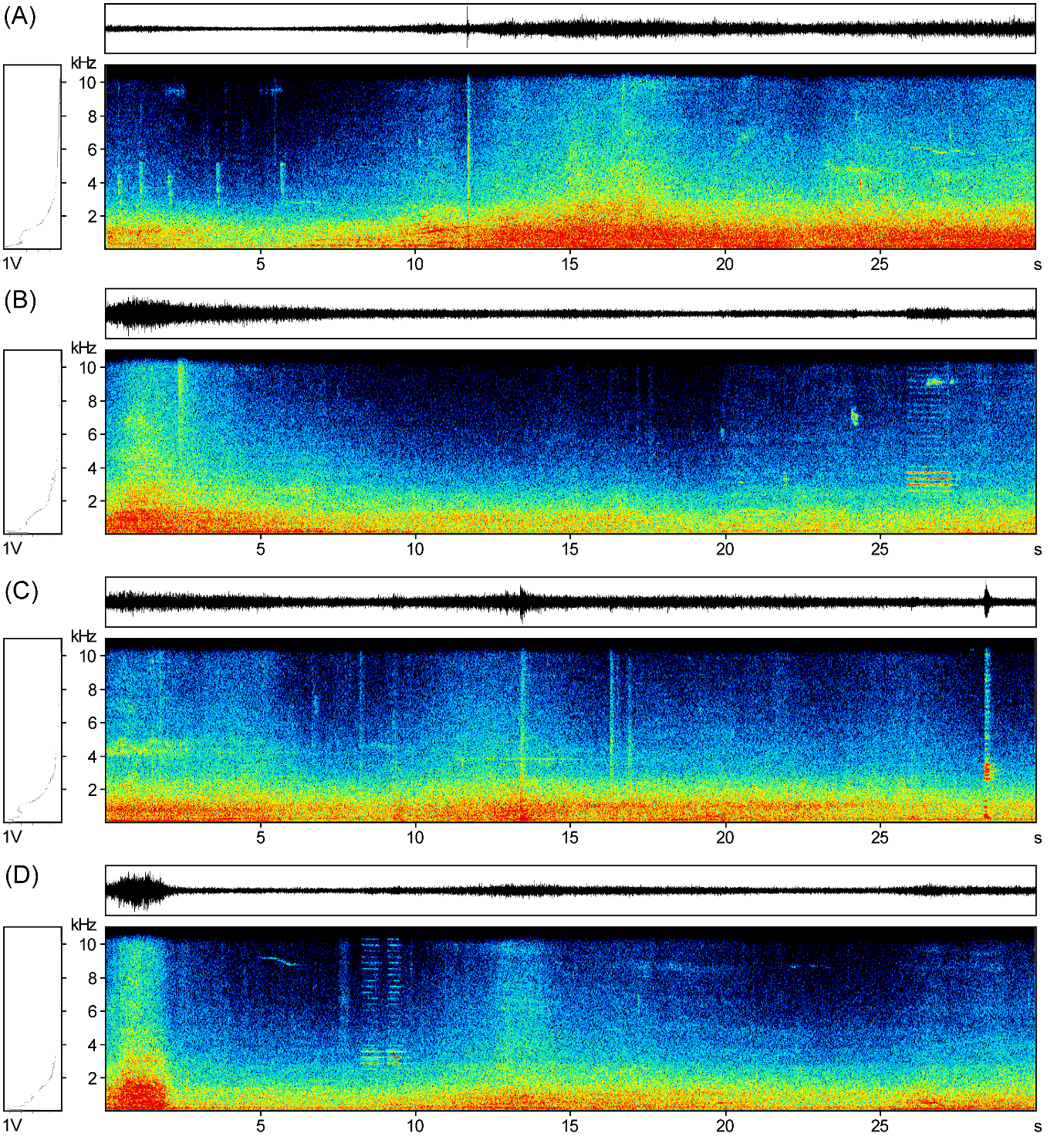

**Figure S8. The exampless of different types of noise.** (A)-(C) Car noise; (D) Motorcycle noise. Oscillograms (above), power spectrum (left) and spectrograms (below).

**Table S9** Assessment of model fit of the discriminant function analyses on the parameters of the best model in three bird species.

| Species | Discriminant  function | Eigenvalue | Percentage variance | Test of function | Wilks’s lambda | Chi-square | *df* | *P* |
| --- | --- | --- | --- | --- | --- | --- | --- | --- |
| Hooded crows | 1 | 0.233 | 95.3 | 1-4 | 0.802 | 117.917 | 16 | < 0.001 |
|  | 2 | 0.009 | 3.8 | 2-4 | 0.989 | 6.068 | 9 | 0.733 |
|  | 3 | 0.002 | 0.7 | 3-4 | 0.998 | 1.082 | 4 | 0.897 |
|  | 4 | 0.001 | 0.2 | 4 | 1.000 | 0.001 | 1 | 0.981 |
| Rose-ringed parakeets | 1 | 0.241 | 85.9 | 1-3 | 0.775 | 136.174 | 15 | < 0.001 |
|  | 2 | 0.032 | 11.1 | 2-3 | 0.962 | 20.846 | 8 | 0.008 |
|  | 3 | 0.008 | 3.0 | 3 | 0.992 | 4.268 | 3 | 0.234 |
| Gracefull prinias | 1 | 0.065 | 77.6 | 1-4 | 0.922 | 43.441 | 20 | 0.002 |
|  | 2 | 0.012 | 14.5 | 2-4 | 0.982 | 9.918 | 12 | 0.623 |
|  | 3 | 0.007 | 7.8 | 3-4 | 0.993 | 3.494 | 6 | 0.745 |
|  | 4 | 0.001 | 0.1 | 4 | 1.000 | 0.010 | 2 | 0.995 |

**Table S10** Akaike’s information criterion (AICc) model comparison results for activity and activity variability in all species and each bird species.

| Species | Model description | *df* | logLik | AICc | ∆AICc | *wi* |
| --- | --- | --- | --- | --- | --- | --- |
| **All species** | **Activity (Number of syllables/day)** ~ |  |  |  |  |  |
|  | Bird, Lock, Noise, Site, Human, Temperature, Bird*Lock, Count*Lock, Lock*Noise, Lock*Site, Lock*Human | 16 | -456966 | 913964.7 | 0.00 | 1 |
|  | Bird, Lock, Noise, Site, Human, Temperature, Bird*Lock, Count*Lock, Lock*Site, Lock*Human | 15 | -457001 | 914033.2 | 68.52 | 0 |
| **All species** | **Activity Variability (CV of the number of syllables/day) ~** |  |  |  |  |  |
|  | Bird, Lock, Noise, Site, Human, Temperature, Bird*Lock, Lock*Noise, Lock*Human | 14 | 88.93 | -149.58 | 0.00 | 0.29 |
|  | Bird, Lock, Noise, Site, Human, Temperature, Bird*Lock, Count*Lock, Lock*Noise, Lock*Human | 15 | 89.59 | -148.85 | 0.73 | 0.20 |
|  | Bird, Lock, Noise, Site, Human, Temperature, Bird*Lock, Lock*Noise | 13 | 87.51 | -148.77 | 0.81 | 0.19 |
|  | Bird, Lock, Noise, Site, Human, Temperature, Bird*Lock, Count*Lock, Lock*Noise | 14 | 88.17 | -148.04 | 1.54 | 0.13 |
|  | Bird, Lock, Noise, Site, Temperature, Bird*Lock, Lock*Noise | 12 | 85.15 | -146.10 | 3.48 | 0.05 |
|  | Bird, Lock, Noise, Site, Human, Temperature, Bird*Lock, Lock*Noise, Lock*Site | 15 | 88.21 | -146.09 | 3.49 | 0.05 |
|  | Bird, Lock, Noise, Site, Human, Temperature, Bird*Lock, Count*Lock, Lock*Noise, Lock*Site, Lock*Human | 16 | 89.12 | -145.87 | 3.71 | 0.05 |
|  | Bird, Lock, Noise, Site, Temperature, Bird*Lock, Count*Lock, Lock*Noise | 13 | 85.92 | -145.59 | 3.99 | 0.04 |
| **Hooded crow** | **Activity (Number of syllables/day)** ~ |  |  |  |  |  |
|  | Lock, Noise, Site, Human, Count*Lock, Lock*Noise, Lock*Site, Lock*Human | 13 | -25425.3 | 50877.20 | 0.00 | 0.70 |
|  | Lock, Noise, Site, Human, Temperature, Count*Lock, Lock*Noise, Lock*Site, Lock*Human | 14 | -25425 | 50878.86 | 1.67 | 0.30 |
| **Hooded crow** | **Activity Variability (CV of the number of syllables/day) ~** |  |  |  |  |  |
|  | Lock, Noise, Site, Human, Temperature, Lock*Noise, Lock*Human | 12 | 184.99 | -345.40 | 0.00 | 0.16 |
|  | Lock, Noise, Site, Human, Lock*Noise, Lock*Human | 11 | 183.92 | -345.34 | 0.06 | 0.16 |
|  | Lock, Noise, Site, Human, Lock*Noise, Lock*Site, Lock*Human | 13 | 185.94 | -345.18 | 0.21 | 0.14 |
|  | Lock, Noise, Site, Human, Temperature, Lock*Noise, Lock*Site, Lock*Human | 14 | 186.93 | -345.05 | 0.35 | 0.13 |
|  | Lock, Noise, Site, Human, Count*Lock, Lock*Noise, Lock*Site, Lock*Human | 14 | 186.80 | -344.80 | 0.60 | 0.12 |
|  | Lock, Noise, Site, Human, Count*Lock, Lock*Noise, Lock*Human | 12 | 184.29 | -343.99 | 1.40 | 0.08 |
|  | Lock, Noise, Site, Human, Temperature, Count*Lock, Lock*Noise, Lock*Site, Lock*Human | 15 | 187.23 | -343.53 | 1.86 | 0.06 |
|  | Lock, Noise, Site, Human, Temperature, Count*Lock, Lock*Noise, Lock*Human | 13 | 185.03 | -343.37 | 2.02 | 0.06 |
|  | Lock, Noise, Site, Human, Count*Lock, Lock*Site, Lock*Human | 13 | 184.63 | -342.56 | 2.84 | 0.04 |
|  | Lock, Noise, Site, Human, Lock*Site, Lock*Human | 12 | 183.15 | -341.71 | 3.69 | 0.03 |
|  | Lock, Noise, Site, Human, Temperature, Lock*Site, Lock*Human | 13 | 184.07 | -341.45 | 3.94 | 0.02 |
| **Rose-ringed**  **parakeet** | **Activity (Number of syllables/day)** ~ |  |  |  |  |  |
|  | Lock, Noise, Site, Human, Temperature, Count*Lock, Lock*Noise, Lock*Site, Lock*Human | 14 | -27201.3 | 54431.34 | 0.00 | 1 |
|  | Lock, Noise, Site, Human, Count*Lock, Lock*Noise, Lock*Site, Lock*Human | 13 | -27235.3 | 54497.28 | 65.94 | 0 |
| **Rose-ringed**  **parakeet** | **Activity Variability (CV of the number of syllables/day) ~** |  |  |  |  |  |
|  | Lock, Noise, Site, Human, Temperature, Count*Lock, Lock*Site, Lock*Human | 14 | 339.72 | -650.62 | 0.00 | 0.41 |
|  | Lock, Noise, Site, Human, Count*Lock, Lock*Site, Lock*Human | 13 | 338.39 | -650.06 | 0.56 | 0.31 |
|  | Lock, Noise, Site, Human, Temperature, Count*Lock, Lock*Noise, Lock*Site, Lock*Human | 15 | 339.87 | -648.81 | 1.81 | 0.17 |
|  | Lock, Noise, Site, Human, Count*Lock, Lock*Noise, Lock*Site, Lock*Human | 14 | 338.46 | -648.11 | 2.51 | 0.12 |
| **Graceful prinia** | **Activity (Number of syllables/day)** ~ |  |  |  |  |  |
|  | Lock, Noise, Site, Human, Temperature, Count*Lock, Lock*Noise, Lock*Site, Lock*Human | 14 | -527547.1 | 1055123 | 0.00 | 1 |
|  | Lock, Noise, Site, Human, Temperature, Count*Lock, Lock*Site, Lock*Human | 13 | -527582.4 | 1055191 | 68.52 | 0 |
| **Graceful prinia** | **Activity Variability (CV of the number of syllables/day) ~** |  |  |  |  |  |
|  | Lock, Noise, Site, Temperature | 9 | -217.52 | 453.49 | 0.00 | 0.09 |
|  | Lock, Noise, Site, Temperature, Count*Lock, Lock*Site | 12 | -214.43 | 453.64 | 0.15 | 0.08 |
|  | Lock, Noise, Site, Temperature, Count*Lock | 10 | -216.62 | 453.80 | 0.31 | 0.08 |
|  | Lock, Noise, Site, Temperature, Lock*Site | 11 | -215.60 | 453.87 | 0.37 | 0.08 |
|  | Lock, Site, Temperature, Count*Lock, Lock*Site | 11 | -215.69 | 454.05 | 0.56 | 0.07 |
|  | Lock, Site, Temperature, Count*Lock | 9 | -217.91 | 454.26 | 0.77 | 0.06 |
|  | Lock, Site, Temperature | 8 | -219.15 | 454.67 | 1.17 | 0.05 |
|  | Lock, Site, Temperature, Lock*Site | 10 | -217.23 | 455.01 | 1.51 | 0.04 |
|  | Lock, Noise, Site, Temperature, Count*Lock, Lock*Noise, Lock*Site | 13 | -214.11 | 455.14 | 1.65 | 0.04 |
|  | Lock, Noise, Site, Human, Temperature, Count*Lock, Lock*Site | 13 | -214.26 | 455.44 | 1.95 | 0.03 |
|  | Lock, Noise, Site, Human, Temperature | 10 | -217.51 | 455.58 | 2.08 | 0.03 |
|  | Lock, Noise, Site, Temperature, Lock*Noise | 10 | -217.52 | 455.59 | 2.09 | 0.03 |
|  | Lock, Site, Human, Temperature, Count*Lock, Lock*Site | 12 | -215.46 | 455.70 | 2.21 | 0.03 |
|  | Lock, Noise, Site, Temperature, Lock*Noise, Lock*Site | 12 | -215.47 | 455.72 | 2.22 | 0.03 |
|  | Noise, Site, Temperature | 8 | -219.71 | 455.78 | 2.28 | 0.03 |
|  | Lock, Noise, Site, Human, Temperature, Lock*Site | 12 | -215.50 | 455.78 | 2.28 | 0.03 |
|  | Lock, Noise, Site, Human, Temperature, Count*Lock | 11 | -216.60 | 455.86 | 2.37 | 0.03 |
|  | Lock, Noise, Site, Temperature, Count*Lock, Lock*Noise | 11 | -216.60 | 455.87 | 2.38 | 0.03 |
|  | Lock, Site, Human, Temperature, Count*Lock | 10 | -217.86 | 456.28 | 2.78 | 0.02 |
|  | Lock, Site, Human, Temperature | 9 | -219.13 | 456.72 | 3.22 | 0.02 |
|  | Lock, Site, Human, Lock*Site | 11 | -217.07 | 456.80 | 3.30 | 0.02 |
|  | Lock, Noise, Site, Human, Temperature, Count*Lock, Lock*Noise, Lock*Site | 14 | -213.88 | 456.82 | 3.33 | 0.02 |
|  | Site | 6 | -222.37 | 456.95 | 3.46 | 0.02 |
|  | Lock, Noise, Site, Human, Temperature, Count*Lock, Lock*Site, Lock*Human | 14 | -214.02 | 457.11 | 3.62 | 0.01 |
|  | Lock, Site, Human, Temperature, Count*Lock, Lock*Site, Lock*Human | 13 | -215.21 | 457.33 | 3.84 | 0.01 |
|  | Noise, Site | 7 | -221.53 | 457.35 | 3.85 | 0.01 |

Bird: Bird species. Lock: Lockdown status. Site: Site category. Human: Human activity. Count: Count down. AICc: Akaike’s information criterion corrected for small sample size. ΔAICc: the difference between the alternative model and best-fitting model. Models are ranked based on the AICc values from the best to the worst model. *: Interaction effect.

**Table 11** Effects of predictor variables on birds’ activity based on generalized and general linear mixed models (GLMM and LMM). Estimates were calculated in % per day for the following units: Temperature – per degree, Noise – per dB, Human activity – per 1 talking event, Lockdown related parameter – per existence of the lockdown (yes/no).

| Species | Dependent  variable | Predictors | Estimate | *z* | *p* | 95% CI | Percent |
| --- | --- | --- | --- | --- | --- | --- | --- |
| All  species | **^#^Activity-**  Number of  syllables/day | (Intercept) | 6.706 | 18.210 | < 0.001 | - | - |
|  |  | Bird_species | 0.131 | **58.938** | **< 0.001** | - | 13.997 |
|  |  | Lockdown_status | 0.270 | **2.048** | **0.041** | - | 30.996 |
|  |  | Human_activity | 0.0001 | **15.114** | **< 0.001** | - | 0.010 |
|  |  | Noise | -0.088 | **-79.861** | **< 0.001** | - | 8.424 |
|  |  | Temperature | 0.187 | **18.949** | **< 0.001** | - | 20.768 |
|  |  | Site_category_residence | -0.946 | **-2.734** | **0.006** | - | 62.394 |
|  |  | Site_category_road | -1.155 | **-3.439** | **0.001** | - | 70.549 |
|  |  | Lockdown_status*Count_down | -0.016 | **-17.871** | **< 0.001** | - | 1.651 |
|  |  | Lockdown_status*Site_category_residence | -0.152 | **-22.224** | **< 0.001** | - | 14.806 |
|  |  | Lockdown_status*Site_category_road | -0.366 | **-42.070** | **< 0.001** | - | 32.489 |
|  |  | Lockdown_status*Noise | -0.009 | **-8.416** | **< 0.001** | - | 0.959 |
|  |  | Lockdown_status*Human_activity | 0.0002 | **24.963** | **< 0.001** | - | 0.022 |
|  |  | Bird_species*Lockdown_status | 0.386 | 146.072 | **< 0.001** | - | 51.348 |
|  | **^#^Activity**  **Variability-**  CV of the  number of  syllables/day | (Intercept) | -0.808 | 4.625 | < 0.001 | -1.164, -0.418 | - |
|  |  | Bird_species | 0.212 | 15.553 | **< 0.001** | **0.185, 0.239** | - |
|  |  | Lockdown_status | 0.579 | 4.014 | **< 0.001** | **0.223, 0.892** | - |
|  |  | Noise | 0.025 | 7.884 | **< 0.001** | **0.018, 0.031** | - |
|  |  | Site_category_residence | 0.183 | 2.520 | **0.012** | **0.036, 0.325** | - |
|  |  | Site_category_road | 0.247 | 3.492 | **< 0.001** | **0.107, 0.387** | - |
|  |  | Human_activity | -0.00003 | 0.658 | 0.510 | -0.0001, 0.00005 | - |
|  |  | Temperature | -0.011 | 3.313 | **0.001** | **-0.017, -0.004** | - |
|  |  | Bird_species*Lockdown_status | -0.061 | 3.768 | **< 0.001** | **-0.093, -0.029** | - |
|  |  | Lockdown_status*Noise | -0.008 | 3.030 | **0.002** | **-0.014, -0.003** | - |
|  |  | Lockdown_status*Human_activity | -0.00007 | 1.657 | 0.097 | -0.0002, 0.00001 | - |
|  |  | Count_down*Lockdown_status | -0.001 | 1.187 | 0.235 | -0.003, 0.0007 | - |
|  |  | Lockdown_status*Site_category_residence | 0.026 | 0.633 | 0.527 | -0.051, 0.113 | - |
| Hooded crow | **Activity-**  Number of  syllables/day | (Intercept) | 9.746 | 22.573 | < 0.001 | 8.900, 10.592 | - |
|  |  | Lockdown_status | -2.293 | 11.906 | **< 0.001** | **-2.670, -1.916** | 89.904 |
|  |  | Noise | -0.062 | 32.300 | **< 0.001** | **-0.066, -0.058** | 6.012 |
|  |  | Site_category_residence | -1.057 | 2.774 | **0.006** | **-1.803, -0.310** | 65.250 |
|  |  | Site_category_road | -0.726 | 1.963 | 0.050 | -1.451, -0.001 | 51.616 |
|  |  | Human_activity | 0.00007 | 5.338 | **< 0.001** | **0.00004, 0.0001** | 0.007 |
|  |  | Count_down*Lockdown_status | 0.055 | 34.878 | **< 0.001** | **0.052, 0.058** | 5.767 |
|  |  | Lockdown_status*Noise | 0.022 | 12.732 | **< 0.001** | **0.0185, 0.025** | 2.291 |
|  |  | Lockdown_status*Site_category_residence | 0.067 | 4.873 | **< 0.001** | **0.040, 0.094** | 7.207 |
|  |  | Lockdown_status*Site_category_road | 0.235 | 16.330 | **< 0.001** | **0.207, 0.263** | 27.815 |
|  |  | Lockdown_status*Human_activity | 0.00018 | 11.500 | **< 0.001** | **0.0001, 0.0002** | 0.019 |
|  |  | Temperature | 0.013 | 0.700 | 0.484 | -0.023, 0.048 | 1.400 |
|  | **^#^Activity**  **Variability-**  CV of the  number of  syllables/day | (Intercept) | -1.248 | 4.511 | < 0.001 | -1.778, -0.688 | - |
|  |  | Lockdown_status | 0.748 | 2.591 | **0.010** | **0.161, 1.325** | - |
|  |  | Noise | 0.033 | 6.152 | **< 0.001** | **0.022, 0.043** | - |
|  |  | Site_category_residence | 0.282 | 2.726 | **0.006** | **0.076, 0.484** | - |
|  |  | Site_category_road | 0.245 | 2.456 | **0.014** | **0.049, 0.440** | - |
|  |  | Human_activity | 0.00006 | 1.370 | 0.171 | -0.00003, 0.0002 | - |
|  |  | Temperature | -0.005 | 1.307 | 0.191 | -0.012, 0.003 | - |
|  |  | Lockdown_status*Noise | -0.013 | 2.634 | **0.008** | **-0.022, -0.003** | - |
|  |  | Lockdown_status*Human_activity | -0.00017 | 3.209 | **0.001** | **-0.0003, -0.00007** | - |
|  |  | Lockdown_status*Site_category_residence | 0.059 | 1.143 | 0.253 | -0.041, 0.166 | - |
|  |  | Lockdown_status*Site_category_road | -0.060 | 1.184 | 0.237 | -0.160, 0.041 | - |
|  |  | Count_down*Lockdown_status | -0.001 | 0.907 | 0.364 | -0.003, 0.001 | - |
| Rose-ringed parakeet | **Activity-**  Number of  syllables/day | (Intercept) | 6.241 | 11.779 | < 0.001 | - | - |
|  |  | Lockdown_status | 0.831 | 4.833 | **< 0.001** | - | 129.561 |
|  |  | Human_activity | -0.0004 | -25.045 | **< 0.001** | - | 0.040 |
|  |  | Noise | -0.044 | -21.666 | **< 0.001** | - | 4.305 |
|  |  | Temperature | 0.124 | 8.265 | **< 0.001** | - | 13.202 |
|  |  | Site_category_residence | -0.519 | -1.066 | 0.287 | - | 40.893 |
|  |  | Site_category_road | -1.614 | -3.415 | **0.001** | - | 81.693 |
|  |  | Lockdown_status*Count_down | -0.026 | -13.760 | **< 0.001** | - | 2.643 |
|  |  | Lockdown_status*Site_category_residence | 0.441 | 28.845 | **< 0.001** | - | 57.643 |
|  |  | Lockdown_status*Site_category_road | 0.571 | 36.727 | **< 0.001** | - | 80.854 |
|  |  | Lockdown_status*Noise | -0.026 | -14.157 | **< 0.001** | - | 2.720 |
|  |  | Lockdown_status*Human_activity | 0.001 | 53.144 | **< 0.001** | - | 0.107 |
|  | **^#^Activity**  **Variability-**  CV of the  number of  syllables/day | (Intercept) | -0.353 | 2.031 | 0.042 | -0.695, -0.009 | - |
|  |  | Lockdown_status | 0.287 | 2.797 | **0.005** | **0.079, 0.506** | - |
|  |  | Noise | 0.019 | 6.739 | **< 0.001** | **0.014, 0.026** | - |
|  |  | Site_category_residence | 0.117 | 1.219 | 0.223 | -0.073, 0.304 | - |
|  |  | Site_category_road | 0.256 | 2.759 | **0.006** | **0.070, 0.437** | - |
|  |  | Human_activity | 0.000 | 1.103 | 0.270 | -0.00004, 0.0001 | - |
|  |  | Temperature | -0.006 | 1.657 | 0.098 | -0.014, 0.001 | - |
|  |  | Count_down*Lockdown_status | -0.004 | 4.140 | **< 0.001** | **-0.006, -0.002** | - |
|  |  | Lockdown_status*Site_category_residence | -0.052 | 1.499 | 0.134 | -0.121, 0.017 | - |
|  |  | Lockdown_status*Site_category_road | -0.121 | 3.470 | **0.001** | **-0.189, -0.052** | - |
|  |  | Lockdown_status*Human_activity | 0.000 | 3.596 | **< 0.001** | **-0.0002, -0.00006** | - |
|  |  | Lockdown_status*Noise | -0.002 | 0.507 | 0.612 | -0.009, 0.005 | - |
| Graceful prinia | **Activity-**  Number of  syllables/day | (Intercept) | 6.972 | 18.936 | < 0.001 | - | - |
|  |  | Lockdown_status | 1.123 | 8.513 | **< 0.001** | - | 207.406 |
|  |  | Human_activity | 0.0001 | 15.114 | **< 0.001** | - | 0.010 |
|  |  | Noise | -0.088 | -79.861 | **< 0.001** | - | 8.424 |
|  |  | Temperature | 0.187 | 18.949 | **< 0.001** | - | 20.563 |
|  |  | Site_category_residence | -0.946 | -2.734 | **0.006** | - | 61.783 |
|  |  | Site_category_road | -1.155 | -3.439 | **0.001** | - | 69.864 |
|  |  | Lockdown_status*Count_down | -0.016 | -17.871 | **< 0.001** | - | 1.635 |
|  |  | Lockdown_status*Site_category_residence | -0.152 | -22.224 | **< 0.001** | - | 14.665 |
|  |  | Lockdown_status*Site_category_road | -0.366 | -42.070 | **< 0.001** | - | 32.182 |
|  |  | Lockdown_status*Noise | -0.009 | -8.416 | **< 0.001** | - | 0.950 |
|  |  | Lockdown_status*Human_activity | 0.0002 | 24.963 | **< 0.001** | - | 0.021 |
|  | **^#^Activity**  **Variability-**  CV of the  number of  syllables/day | (Intercept) | 1.172 | 2.048 | 0.041 | 0.0127, 2.247 | - |
|  |  | Lockdown_status | -0.190 | 0.677 | 0.498 | -0.751, 0.407 | - |
|  |  | Noise | 0.015 | 1.582 | 0.114 | -0.004, 0.035 | - |
|  |  | Site_category_residence | 0.191 | 0.930 | 0.353 | -0.210, 0.594 | - |
|  |  | Site_category_road | 0.531 | 2.495 | **0.013** | **0.117, 0.953** | - |
|  |  | Temperature | -0.029 | 2.528 | **0.012** | **-0.051, -0.006** | - |
|  |  | Count_down*Lockdown_status | 0.005 | 1.544 | 0.123 | -0.002, 0.011 | - |
|  |  | Lockdown_status*Site_category_residence | 0.147 | 1.278 | 0.201 | -0.079, 0.374 | - |
|  |  | Lockdown_status*Site_category_road | 0.260 | 2.044 | **0.041** | **0.011, 0.512** | - |
|  |  | Lockdown_status*Noise | -0.006 | 0.487 | 0.627 | -0.029, 0.018 | - |
|  |  | Human_activity | -0.00005 | 0.464 | 0.642 | -0.0003, 0.0002 | - |
|  |  | Lockdown_status*Human_activity | 0.0001 | 0.698 | 0.485 | -0.0003, 0.0004 | - |

CV: coefficient of variance. *: Interaction effect. 95% confidence intervals of the parameters that did not overlap zero are indicated in bold. **#**: model average.

**Table S12** Results from post hoc tests comparing the differences in activity and activity variability between no lockdown period and lockdown period in all species for each site. Estimates were calculated in % per day for the following units: Temp – per degree, Noise – per dB, Human activity – per 1 Human_activitying event, Lockdown related parameter – per existence of the lockdown (yes/no).

| Species | Dependent variable | Site category | Estimate | *P* | Percent |
| --- | --- | --- | --- | --- | --- |
| All species | **Activity**-  Number of syllables/day | Road | -0.073 | 0.767 | 7.040 |
|  |  | Residence | -0.286 | 0.244 | 24.874 |
|  |  | Park | -0.438 | 0.074 | 35.467 |
|  | **Activity variability**-  CV of the number of syllables/day | Road | -0.037 | 0.260 | - |
|  |  | Residence | -0.016 | 0.663 | - |
|  |  | Park | -0.0006 | 0.988 | - |
| Hooded crow | **Activity**-  Number of syllables/day | Road | 0.155 | 0.659 | 16.766 |
|  |  | Residence | 0.323 | 0.359 | 38.127 |
|  |  | Park | 0.390 | 0.268 | 47.698 |
|  | **Activity variability**-  CV of the number of syllables/day | Road | -0.046 | 0.217 | - |
|  |  | Residence | -0.078 | 0.080 | - |
|  |  | Park | -0.034 | 0.461 | - |
| Rose-ringed parakeet | **Activity**-  Number of events/day | Road | -0.209 | 0.138 | 18.860 |
|  |  | Residence | -0.079 | 0.577 | 7.596 |
|  |  | Park | 0.362 | **0.010** | 43.620 |
|  | **Activity variability**-  CV of the number of syllables/day | Road | -0.029 | 0.332 | - |
|  |  | Residence | -0.070 | **0.038** | - |
|  |  | Park | -0.128 | **0.0003** | - |
| Graceful prinia | **Activity-**  Number of syllables/day | Road | -0.464 | 0.302 | 37.124 |
|  |  | Residence | -1.838 | **<0.0001** | 84.086 |
|  |  | Park | -1.922 | **<0.0001** | 85.369 |
|  | **Activity variability**-  CV of the number of syllables/day | Road | -0.038 | 0.781 | - |
|  |  | Residence | 0.142 | 0.285 | - |
|  |  | Park | 0.303 | **0.030** | - |

CV: coefficient of variance

**Table S16** Effects of predictor variables on birds’ activity based on generalized and general linear mixed models (GLMM and LMM). Estimates were calculated in % per day for the following units: Temperature – per degree, Noise – per dB, Human activity – per 1 talking event, Lockdown related parameter – per existence of the lockdown (yes/no).

| Species | Dependent  variable | Predictors | Estimate | *z* | *t* | *p* | 95% CI | Percent |
| --- | --- | --- | --- | --- | --- | --- | --- | --- |
| All species | **Activity-**  Number of  events/day | (Intercept) | 7.042 | 18.271 | - | < 0.001 | - | - |
|  |  | Bird_species | -0.642 | -76.465 | **-** | **< 0.001** | - | 47.38 |
|  |  | Lockdown_status | 0.209 | 1.259 | - | 0.208 | - | 23.24 |
|  |  | Human_activity | -0.0001 | -3.351 | **-** | **0.001** | **-** | 0.01 |
|  |  | Noise | -0.049 | -14.612 | **-** | **< 0.001** | - | 4.78 |
|  |  | Temperature | 0.052 | 3.975 | **-** | **< 0.001** | **-** | 5.34 |
|  |  | Site_category_residence | -0.722 | -2.739 | **-** | **0.006** | **-** | 51.42 |
|  |  | Site_category_road | -0.920 | -3.596 | **-** | **< 0.001** | **-** | 60.15 |
|  |  | Lockdown_status*Count_down | 0.008 | 3.214 | **-** | **0.001** | **-** | 0.80 |
|  |  | Lockdown_status*Site_category_residence | -0.092 | -3.565 | **-** | **< 0.001** | **-** | 8.79 |
|  |  | Lockdown_status*Site_category_road | 0.039 | 1.495 | - | 0.135 | **-** | 3.98 |
|  |  | Lockdown_status*Noise | -0.021 | -7.266 | **-** | **< 0.001** | **-** | 2.08 |
|  |  | Lockdown_status*Human_activity | 0.0004 | 14.956 | **-** | **< 0.001** | **-** | 0.04 |
|  |  | Bird_species*Lockdown_status | 0.269 | 27.540 | **-** | **< 0.001** | **-** | 30.87 |
|  | **^#^Activity**  **Variability-**  CV of the  number of  events/day | (Intercept) | -0.689 | - | -4.312 | < 0.001 | **-**1.0171, -0.2238 | - |
|  |  | Bird_species | 0.210 | - | 15.391 | **< 0.001** | **0.1815, 0.2372** | - |
|  |  | Lockdown_status | 0.378 | - | 2.898 | **0.004** | **-0.0542, 0.6611** | - |
|  |  | Human_activity | -0.0001 | - | -2.021 | **0.043** | **-0.0001, 0.00002** | - |
|  |  | Noise | 0.023 | - | 7.956 | **< 0.001** | **0.0151, 0.0288** | - |
|  |  | Temperature | -0.017 | - | -4.990 | **< 0.001** | **-0.026, -0.0096** | - |
|  |  | Site_category_residence | 0.170 | - | 2.485 | **0.023** | **0.0184, 0.3051** | - |
|  |  | Site_category_road | 0.227 | - | 3.389 | **0.003** | **0.0763, 0.3573** | - |
|  |  | Lockdown_status*Noise | -0.005 | - | -2.094 | **0.036** | **-0.0117, 0.0003** | - |
|  |  | Bird_species*Lockdown_status | -0.056 | - | -3.497 | **< 0.001** | **-0.0866, -0.0237** | - |
| Hooded  crow | #Activity-  Number of  events/day | (Intercept) | 6.446 | 19.191 | - | < 0.001 | 5.7095, 7.3195 | - |
|  |  | Lockdown_type | 0.007 | 0.033 | - | 0.974 | -0.4547, 0.4164 | 0.70 |
|  |  | Human_activity | -0.0001 | -4.264 | **-** | **< 0.001** | **-0.0002, -0.0001** | 0.01 |
|  |  | Noise | -0.027 | -6.061 | **-** | **< 0.001** | **-0.0354, -0.0178** | 2.66 |
|  |  | Site_category_residence | -0.746 | -2.247 | **-** | **0.025** | **-1.3985, -0.0938** | 52.57 |
|  |  | Site_category_road | -0.765 | -2.375 | **-** | **0.018** | **-1.3971, -0.1315** | 53.47 |
|  |  | Lockdown_status*Count_down | 0.013 | 5.209 | **-** | **< 0.001** | **0.0082, 0.0192** | 1.31 |
|  |  | Lockdown_status*Site_category_residence | -0.089 | -2.477 | **-** | **0.013** | **-0.1596, -0.0166** | 8.52 |
|  |  | Lockdown_status*Site_category_road | 0.158 | 4.556 | **-** | **< 0.001** | **0.0890, 0.2275** | 17.12 |
|  |  | Lockdown_status*Noise | -0.012 | -3.221 | **-** | **0.001** | **-0.0194, -0.0046** | 1.19 |
|  |  | Lockdown_status*Human_activity | 0.0003 | 8.887 | **-** | **< 0.001** | **0.0003, 0.0004** | 0.03 |
|  | **^#^Activity**  **Variability-**  CV of the  number of  events/day | (Intercept) | -1.181 | - | -6.232 | < 0.001 | -1.649, -0.639 | - |
|  |  | Lockdown_status | 0.694 | - | 4.741 | **< 0.001** | **0.163, 1.068** | - |
|  |  | Human_activity | 0.0000 | - | 0.721 | 0.471 | -0.0001, 0.0001 | - |
|  |  | Noise | 0.032 |  | 9.356 | **< 0.001** | **0.022, 0.039** | - |
|  |  | Temperature | -0.007 | - | -1.795 | 0.073 | -0.017, 0.002 | - |
|  |  | Site_category_residence | 0.306 | - | 3.157 | **0.006** | **0.071, 0.488** | - |
|  |  | Site_category_road | 0.235 | - | 2.497 | **0.023** | **0.038, 0.422** | - |
|  |  | Lockdown_status*Noise | -0.012 | - | -4.005 | **< 0.001** | **-0.018, -0.004** | - |
|  |  | Lockdown_status*Human_activity | -0.0001 | - | -2.378 | **0.018** | **-0.0002, -0.00001** | - |
| Rose-ringed  parakeet | **^#^Activity-**  Number of  events/day | (Intercept) | 5.058 | 9.767 | - | < 0.001 | - | - |
|  |  | Lockdown_status | 0.276 | 1.051 | - | 0.293 | - | 31.78 |
|  |  | Human_activity | -0.0003 | -6.814 | **-** | **< 0.001** | **-** | 0.03 |
|  |  | Noise | -0.050 | -8.109 | **-** | **< 0.001** | - | 4.88 |
|  |  | Temperature | 0.090 | 6.584 | **-** | **< 0.001** | **-** | 9.42 |
|  |  | Site_category_residence | -0.479 | -1.146 | - | 0.252 | - | 38.06 |
|  |  | Site_category_road | -1.172 | -2.889 | **-** | **0.004** | - | 69.03 |
|  |  | Lockdown_status*Site_category_residence | 0.390 | 8.039 | **-** | **< 0.001** | **-** | 47.70 |
|  |  | Lockdown_status*Site_category_road | 0.423 | 8.782 | **-** | **< 0.001** | **-** | 52.65 |
|  |  | Lockdown_status*Noise | -0.022 | -4.276 | **-** | **< 0.001** | **-** | 2.18 |
|  |  | Lockdown_status*Human_activity | 0.001 | 15.119 | **-** | **< 0.001** | **-** | 0.10 |
|  | **^#^Activity**  **Variability-**  CV of the  number of  events/day | (Intercept) | -0.102 | - | -0.670 | 0.504 | -0.3567, 0.3557 | - |
|  |  | Lockdown_status | 0.248 | - | 2.420 | **0.016** | **0.0830, 0.3802** | - |
|  |  | Human_activity | -0.0001 | - | -1.665 | 0.097 | -0.0001, 0.0000, | - |
|  |  | Noise | 0.020 | - | 8.471 | **< 0.001** | **0.0133, 0.023** | - |
|  |  | Temperature | -0.014 | - | -2.988 | **0.004** | **-0.0274, -0.0074** | - |
|  |  | Lockdown_status*Count_down | -0.002 | - | -1.651 | 0.103 | -0.0041, 0.0005 | - |
|  |  | Lockdown_status*Noise | -0.004 | - | -1.777 | 0.076 | -0.0075, 0.0008 | - |
|  |  | Lockdown_status*Human_activity | -0.0001 | - | -1.498 | 0.135 | -0.0001, 0.00001 | - |
| Graceful  prinia | **^#^Activity-**  Number of  events/day | (Intercept) | 1.536 | 1.139 | - | 0.255 | -1.1049, 4.2863 | - |
|  |  | Lockdown_status | 2.201 | 6.528 | **-** | **< 0.001** | **1.2948, 2.9911** | 83.40 |
|  |  | Human_activity | -0.0001 | -1.481 | - | 0.139 | -0.0003, 0.00004 | 0.01 |
|  |  | Noise | -0.148 | -21.105 | **-** | **< 0.001** | **-0.1657, -0.1326** | 13.76 |
|  |  | Temperature | 0.416 | 7.606 | **-** | **< 0.001** | **0.3083, 0.5235** | 51.59 |
|  |  | Site_category_residence | -1.126 | -1.710 | - | 0.087 | -2.4253, 0.1648 | 67.57 |
|  |  | Site_category_road | -1.519 | -2.368 | **-** | **0.018** | **-2.7761, -0.2513** | 78.11 |
|  |  | Lockdown_status*Count_down | -0.077 | -9.716 | **-** | **< 0.001** | **-0.0919, -0.0609** | 7.41 |
|  |  | Lockdown_status*Site_category_residence | -0.163 | -3.044 | **-** | **0.002** | **-0.2748, -0.0410** | 15.04 |
|  |  | Lockdown_status*Site_category_road | -0.909 | -11.494 | **-** | **< 0.001** | **-1.0850, -0.7472** | 59.71 |
|  |  | Lockdown_status*Human_activity | 0.001 | 8.404 | **-** | **< 0.001** | **0.0005, 0.0008** | 0.10 |
|  | **^#^Activity**  **Variability-**  CV of the  number of  events/day | (Intercept) | 1.353 | - | 2.276 | 0.024 | -0.4557, 2.6207 | - |
|  |  | Lockdown_status | -0.482 | - | -3.197 | **0.002** | **-1.0859, 0.4182** | - |
|  |  | Noise | 0.025 | - | 2.800 | **0.006** | **0.0049, 0.0461** | - |
|  |  | Temperature | -0.072 | - | -3.485 | **0.001** | **-0.1125, -0.0119** | - |
|  |  | Site_category_residence | 0.208 | - | 0.990 | 0.331 | -0.1579, 0.6552 | - |
|  |  | Site_category_road | 0.415 | - | 2.000 | 0.055 | 0.0730, 0.9539 | - |
|  |  | Lockdown_status*Count_down | 0.011 | - | 2.399 | **0.017** | **0.0012, 0.0199** | - |
|  |  | Lockdown_status*Site_category_residence | 0.124 | - | 0.976 | 0.330 | -0.1430, 0.3940 | - |
|  |  | Lockdown_status*Site_category_road | 0.308 | - | 2.157 | **0.032** | **0.0249, 0.6052** | - |

CV: coefficient of variance. *: Interaction effect. 95% confidence intervals of the parameters that did not overlap zero are indicated in bold. **#**: model average.

**Table S17** Akaike’s information criterion (AICc) model comparison results for activity and activity variability in all species and each bird species.

| Species | Model description | *df* | logLik | AICc | ∆AICc | *wi* |
| --- | --- | --- | --- | --- | --- | --- |
| **All species** | **Activity (Number of events/day)** ~ |  |  |  |  |  |
|  | Bird, Lock, Noise, Site, Human, Temperature, Bird*Lock, Count*Lock, Lock*Noise, Lock*Site, Lock*Human | 16 | -24379.21 | 48790.83 | 0.00 | 0.98 |
|  | Bird, Lock, Noise, Site, Human, Temperature, Bird*Lock, Noise*Lock, Lock*Site, Lock*Human | 15 | -24384.24 | 48798.84 | 8.01 | 0.02 |
| **All species** | **Activity Variability (CV of the number of events/day) ~** |  |  |  |  |  |
|  | Bird, Lock, Noise, Site, Human, Temperature, Bird*Lock, Lock*Noise | 13 | 215.89 | -405.48 | 0.00 | 0.21 |
|  | Bird, Lock, Noise, Site, Human, Temperature, Bird*Lock, Lock*Noise, Lock*Human | 14 | 216.37 | -404.40 | 1.08 | 0.12 |
|  | Bird, Lock, Noise, Site, Human, Temperature, Bird*Lock, Lock*Noise, Lock*Site | 15 | 217.39 | -404.38 | 1.10 | 0.12 |
|  | Bird, Lock, Noise, Site, Human, Temperature, Bird*Lock, Count*Lock, Lock*Noise | 14 | 215.95 | -403.55 | 1.93 | 0.08 |
|  | Bird, Lock, Noise, Site, Temperature, Bird*Lock, Lock*Noise | 12 | 213.89 | -403.54 | 1.95 | 0.08 |
|  | Bird, Lock, Noise, Site, Human, Temperature, Bird*Lock | 12 | 213.70 | -403.15 | 2.33 | 0.07 |
|  | Bird, Lock, Noise, Site, Human, Temperature, Bird*Lock, Count*Lock, Lock*Noise, Lock*Site | 16 | 217.52 | -402.60 | 2.88 | 0.05 |
|  | Bird, Lock, Noise, Site, Temperature, Bird*Lock | 11 | 212.40 | -402.58 | 2.90 | 0.05 |
|  | Bird, Lock, Noise, Site, Human, Temperature, Bird*Lock, Lock*Site | 14 | 215.43 | -402.53 | 2.95 | 0.05 |
|  | Bird, Lock, Noise, Site, Human, Temperature, Bird*Lock, Count*Lock, Lock*Noise, Lock*Human | 15 | 216.45 | -402.51 | 2.97 | 0.05 |
|  | Bird, Lock, Noise, Site, Human, Temperature, Bird*Lock, Lock*Noise, Lock*Site, Lock*Human | 16 | 217.45 | -402.46 | 3.02 | 0.05 |
|  | Bird, Lock, Noise, Site, Human, Temperature, Bird*Lock, Lock*Human | 13 | 214.26 | -402.22 | 3.26 | 0.04 |
|  | Bird, Lock, Noise, Site, Temperature, Bird*Lock, Count*Lock, Lock*Noise | 13 | 213.94 | -401.58 | 3.90 | 0.03 |
| **Hooded crow** | **Activity (Number of events/day)** ~ |  |  |  |  |  |
|  | Lock, Noise, Site, Human, Count*Lock, Lock*Noise, Lock*Site, Lock*Human | 13 | -4410.91 | 8848.64 | 0.00 | 0.70 |
|  | Lock, Noise, Site, Human, Temperature, Count*Lock, Lock*Noise, Lock*Site, Lock*Human | 14 | -4410.76 | 8850.47 | 1.83 | 0.28 |
|  | Lock, Noise, Site, Human, Count*Lock, Lock*Site, Lock*Human | 12 | -4416.08 | 8856.87 | 8.22 | 0.01 |
|  | Lock, Noise, Site, Human, Temperature, Count*Lock, Lock*Site, Lock*Human | 13 | -4415.58 | 8858.00 | 9.35 | 0.01 |
| **Hooded crow** | **Activity Variability (CV of the number of events/day) ~** |  |  |  |  |  |
|  | Lock, Noise, Site, Human, Temperature, Lock*Noise, Lock*Human | 12 | 213.04 | -401.37 | 0.00 | 0.26 |
|  | Lock, Noise, Site, Human, Lock*Noise, Lock*Human | 11 | 211.47 | -400.35 | 1.03 | 0.16 |
|  | Lock, Noise, Site, Human, Count*Lock, Lock*Noise, Lock*Human | 12 | 212.22 | -399.74 | 1.64 | 0.11 |
|  | Lock, Noise, Site, Human, Temperature, Count*Lock, Lock*Noise, Lock*Human | 13 | 213.06 | -399.30 | 2.08 | 0.09 |
|  | Lock, Noise, Site, Human, Temperature, Lock*Noise, Lock*Site, Lock*Human | 14 | 213.93 | -398.90 | 2.47 | 0.08 |
|  | Lock, Noise, Site, Human, Temperature, Lock*Noise, Lock*Site | 13 | 212.52 | -398.22 | 3.16 | 0.05 |
|  | Lock, Noise, Site, Temperature, Lock*Noise | 10 | 209.26 | -398.03 | 3.35 | 0.05 |
|  | Lock, Noise, Site, Human, Temperature, Lock*Noise | 11 | 210.24 | -397.88 | 3.49 | 0.05 |
|  | Lock, Noise, Site, Human, Lock*Noise, Lock*Site, Lock*Human | 13 | 212.32 | -397.81 | 3.56 | 0.04 |
|  | Lock, Noise, Site, Lock*Noise | 9 | 208.01 | -397.61 | 3.76 | 0.04 |
|  | Lock, Noise, Site, Human, Lock*Noise, Lock*Site | 12 | 211.10 | -397.48 | 3.89 | 0.04 |
|  | Lock, Noise, Human, Temperature, Lock*Noise, Lock*Human | 10 | 208.94 | -397.38 | 3.99 | 0.04 |
| **Rose-ringed**  **parakeet** | **Activity (Number of events/day)** ~ |  |  |  |  |  |
|  | Lock, Noise, Site, Human, Temperature, Lock*Noise, Lock*Site, Lock*Human | 13 | -3029.05 | 6084.94 | 0.00 | 0.74 |
|  | Lock, Noise, Site, Human, Temperature, Count*Lock, Lock*Noise, Lock*Site, Lock*Human | 14 | -3029.05 | 6087.05 | 2.12 | 0.26 |
| **Rose-ringed**  **parakeet** | **Activity Variability (CV of the number of events/day) ~** |  |  |  |  |  |
|  | Lock, Noise, Human, Temperature, Count*Lock, Lock*Noise, Lock*Human | 11 | 393.36 | -764.11 | 0.00 | 0.08 |
|  | Lock, Noise, Human, Temperature, Count*Lock, Lock*Noise | 10 | 392.25 | -763.99 | 0.12 | 0.08 |
|  | Lock, Noise, Human, Temperature, Lock*Noise, Lock*Human | 10 | 392.03 | -763.55 | 0.56 | 0.06 |
|  | Lock, Noise, Human, Temperature, Lock*Human | 9 | 390.85 | -763.30 | 0.81 | 0.06 |
|  | Noise, Human, Temperature | 7 | 388.76 | -763.27 | 0.84 | 0.05 |
|  | Lock, Noise, Site, Human, Temperature, Count*Lock, Lock*Noise, Lock*Human | 13 | 395.03 | -763.22 | 0.89 | 0.05 |
|  | Lock, Noise, Human, Temperature, Count*Lock, Lock*Noise, Lock*Site, Lock*Human | 10 | 391.82 | -763.13 | 0.98 | 0.05 |
|  | Lock, Noise, Site, Human, Temperature, Count*Lock, Lock*Human | 9 | 390.73 | -763.05 | 1.06 | 0.05 |
|  | Lock, Noise, Site, Human, Temperature, Count*Lock, Lock*Noise | 12 | 393.86 | -763.01 | 1.10 | 0.05 |
|  | Lock, Noise, Site, Human, Temperature, Lock*Noise, Lock*Human | 12 | 393.83 | -762.93 | 1.18 | 0.05 |
|  | Lock, Noise, Human, Temperature, Count*Lock | 9 | 390.64 | -762.87 | 1.24 | 0.04 |
|  | Lock, Noise, Site, Human, Temperature, Count*Lock, Lock*Site, Lock*Human | 14 | 395.91 | -762.85 | 1.26 | 0.04 |
|  | Lock, Noise, Human, Temperature | 8 | 389.51 | -762.69 | 1.42 | 0.04 |
|  | Lock, Noise, Site, Human, Temperature, Lock*Human | 11 | 392.64 | -762.67 | 1.44 | 0.04 |
|  | Noise, Site, Human, Temperature | 9 | 390.42 | -762.43 | 1.68 | 0.04 |
|  | Lock, Noise, Site, Human, Temperature, Lock*Site, Lock*Human | 13 | 394.63 | -762.42 | 1.69 | 0.04 |
|  | Lock, Noise, Site, Human, Temperature, Lock*Noise | 11 | 392.47 | -762.33 | 1.78 | 0.03 |
|  | Lock, Noise, Site, Human, Temperature, Count*Lock, Lock*Human | 12 | 393.50 | -762.28 | 1.83 | 0.03 |
|  | Lock, Noise, Site, Human, Temperature | 10 | 391.23 | -761.96 | 2.15 | 0.03 |
|  | Lock, Noise, Site, Human, Temperature, Count*Lock | 11 | 392.26 | -761.91 | 2.20 | 0.03 |
|  | Lock, Noise, Site, Human, Temperature, Count*Lock, Lock*Noise, Lock*Site, Lock*Human | 15 | 396.35 | -761.58 | 2.53 | 0.02 |
|  | Lock, Noise, Site, Human, Temperature, Lock*Noise, Lock*Site, Lock*Human | 14 | 395.04 | -761.10 | 3.01 | 0.02 |
|  | Lock, Noise, Site, Human, Temperature, Count*Lock, Lock*Site | 13 | 393.67 | -760.49 | 3.62 | 0.01 |
| **Graceful prinia** | **Activity (Number of events/day)** ~ |  |  |  |  |  |
|  | Lock, Noise, Site, Human, Temperature, Count*Lock, Lock*Site, Lock*Human | 13 | -6704.26 | 13435.34 | 0.00 | 0.72 |
|  | Lock, Noise, Site, Human, Temperature, Count*Lock, Lock*Noise, Lock*Site, Lock*Human | 14 | -6704.16 | 13437.28 | 1.94 | 0.28 |
| **Graceful prinia** | **Activity Variability (CV of the number of events/day) ~** |  |  |  |  |  |
|  | Lock, Noise, Site, Temperature, Count*Lock, Lock*Site | 12 | -194.55 | 414.03 | 0.00 | 0.25 |
|  | Lock, Noise, Site, Temperature, Count*Lock | 10 | -196.89 | 414.44 | 0.41 | 0.20 |
|  | Lock, Noise, Site, Temperature, Count*Lock, Lock*Noise, Lock*Site | 13 | -194.44 | 415.97 | 1.94 | 0.09 |
|  | Lock, Noise, Site, Human, Temperature, Count*Lock, Lock*Site | 13 | -194.49 | 416.07 | 2.04 | 0.09 |
|  | Lock, Noise, Site, Temperature, Count*Lock, Lock*Noise | 11 | -196.82 | 416.42 | 2.39 | 0.08 |
|  | Lock, Noise, Site, Human, Temperature, Count*Lock | 11 | -196.88 | 416.56 | 2.52 | 0.07 |
|  | Noise, Site, Temperature | 8 | -200.33 | 417.09 | 3.06 | 0.05 |
|  | Lock, Noise, Site, Temperature | 9 | -199.31 | 417.16 | 3.13 | 0.05 |
|  | Lock, Noise, Site, Temperature, Lock*Site | 11 | -197.40 | 417.59 | 3.56 | 0.04 |
|  | Lock, Noise, Site, Human, Temperature, Count*Lock, Lock*Noise, Lock*Site | 14 | -194.35 | 417.97 | 3.94 | 0.03 |
|  | Lock, Noise, Site, Human, Temperature, Count*Lock, Lock*Site, Lock*Human | 14 | -194.38 | 418.03 | 4.00 | 0.03 |

Bird: Bird species. Lock: Lockdown status. Site: Site category. Human: Human activity. Count: Count down. AICc: Akaike’s information criterion corrected for small sample size. ΔAICc: the difference between the alternative model and best-fitting model. Models are ranked based on the AICc values from the best to the worst model. *: Interaction effect.

**Table S18** Effects of predictor variables on birds’ activity based on generalized and general linear mixed models (GLMM and LMM). Estimates were calculated in % per day for the following units: Temperature – per degree, Noise – per dB, Human activity – per 1 talking event, Lockdown related parameter – per existence of the lockdown (yes/no).

| Species | Dependent  variable | Predictors | Estimate | *z* | t | *p* | 95% CI | Percent |
| --- | --- | --- | --- | --- | --- | --- | --- | --- |
| All species | **Activity-**  Number of  events/day | (Intercept) | 7.555 | 22.048 | - | **<** 0.001 | - | - |
|  |  | Bird_species | -0.623 | -84.742 | - | **< 0.001** | - | 46.367 |
|  |  | Lockdown_status | -0.064 | -0.437 | - | 0.662 | - | 6.200 |
|  |  | Human_activity | -0.00002 | -0.725 | - | 0.469 | - | 0.002 |
|  |  | Noise | -0.062 | -21.471 | - | **< 0.001** | - | 6.012 |
|  |  | Temperature | 0.055 | 5.114 | - | **< 0.001** | - | 5.711 |
|  |  | Site_category_residence | -0.762 | -2.927 | - | **0.003** | - | 54.393 |
|  |  | Site_category_road | -0.917 | -3.626 | - | **<0.001** | - | 61.829 |
|  |  | Lockdown_status*Count_down | 0.005 | 2.552 | - | **0.011** | - | 0.521 |
|  |  | Lockdown_status*Site_category_residence | -0.068 | -3.213 | - | **0.001** | - | 6.903 |
|  |  | Lockdown_status*Site_category_road | 0.068 | 3.003 | - | **0.003** | - | 7.459 |
|  |  | Lockdown_status*Noise | -0.014 | -5.400 | - | **< 0.001** | - | 1.488 |
|  |  | Lockdown_status*Human_activity | 0.0004 | 14.825 | - | **< 0.001** | - | 0.043 |
|  |  | Bird_species*Lockdown_status | 0.253 | 29.155 | - | **< 0.001** | - | 31.379 |
|  | **^#^Activity**  **Variability-**  CV of the  number of  events/day | (Intercept) | -0.835 | - | -5.908 | < 0.001 | -1.167, -0.527 | - |
|  |  | Bird_species | 0.208 | - | 16.856 | **< 0.001** | **0.183, 0.232** | - |
|  |  | Lockdown_status | 0.466 | - | 3.946 | **< 0.001** | **0.157, 0.746** | - |
|  |  | Human_activity | -0.00005 | - | -2.224 | **0.026** | **-0.0001, -0.00002** | - |
|  |  | Noise | 0.024 | - | 9.033 | **< 0.001** | **0.018, 0.029** | - |
|  |  | Temperature | -0.012 | - | -4.571 | **< 0.001** | **-0.017, -0.0052** | - |
|  |  | Site_category_residence | 0.175 | - | 2.546 | **0.021** | **0.035, 0.313** | - |
|  |  | Site_category_road | 0.232 | - | 3.471 | **0.003** | **0.093, 0.363** | - |
|  |  | Lockdown_status*Noise | -0.007 | - | -2.908 | **0.004** | **-0.012, -0.002** | - |
|  |  | Bird_species*Lockdown_status | -0.056 | - | -3.838 | **< 0.001** | **-0.085, -0.027** | - |
| Hooded  crow | **^#^Activity-**  Number of  events/day | (Intercept) | 6.901 | 22.934 | - | < 0.001 | 6.165, 7.608 | - |
|  |  | Lockdown_status | -0.627 | -7.950 | - | **< 0.001** | **-0.887, -0.169** | 46.581 |
|  |  | Human_activity | -0.0001 | -3.540 | - | **< 0.001** | **-0.0002, -0.00005** | 0.010 |
|  |  | Noise | -0.036 | -11.648 | - | **< 0.001** | **-0.042, -0.027** | 3.536 |
|  |  | Site_category_residence | -0.756 | -2.294 | - | **0.022** | **-1.399, -0.102** | 53.046 |
|  |  | Site_category_road | -0.773 | -2.413 | - | **0.016** | **-1.408, -0.149** | 54.376 |
|  |  | Count_down*Lockdown_status | 0.015 | 6.235 | - | **< 0.001** | **0.010, 0.020** | 1.542 |
|  |  | Lockdown_status*Site_category_residence | -0.064 | -2.399 | - | **0.017** | **-0.132, -0.015** | 6.385 |
|  |  | Lockdown_status*Site_category_road | 0.193 | 6.934 | - | **< 0.001** | **0.143, 0.261** | 22.140 |
|  |  | Lockdown_status*Human_activity | 0.0003 | 9.935 | - | **< 0.001** | **0.0003, 0.0004** | 0.032 |
|  | **^#^Activity**  **ariability-**  CV of the  number of  events/day | (Intercept) | -1.161 | - | -7.446 | **< 0.001** | **-1.662,-0.882** | - |
|  |  | Lockdown_type | 0.841 | - | 6.346 | **< 0.001** | **0.450, 1.122** | - |
|  |  | Human_activity | 0.00004 | - | 0.999 | 0.318 | -0.00001, 0.0001 | - |
|  |  | Noise | 0.032 | - | 10.567 | **< 0.001** | **0.025, 0.038** | - |
|  |  | Count_down*Lockdown_status | -0.001 | - | -1.143 | 0.256 | **-0.003, 0.001** | - |
|  |  | Lockdown_status*Noise | -0.014 | - | -5.163 | **< 0.001** | **-0.020, -0.007** | - |
|  |  | Lockdown_status*Human_activity | -0.0001 | - | -2.717 | **0.007** | **-0.0002, -0.00003** | - |
| Rose-ringed  parakeet | **^#^Activity-**  Number of  events/day | (Intercept) | 6.152 | 12.169 | - | < 0.001 | 5.206, 7.255 | - |
|  |  | Lockdown_type | -0.051 | -0.213 | - | 0.832 | -0.604, 0.408 | 4.972 |
|  |  | Human_activity | 0.0003 | -6.909 | - | **< 0.001** | **-0.0004, -0.0002** | 0.030 |
|  |  | Noise | -0.063 | -11.621 | - | **< 0.001** | **-0.074, -0.052** | 6.106 |
|  |  | Temperature | 0.068 | 4.838 | - | **< 0.001** | **0.035, 0.094** | 7.037 |
|  |  | Site_category_residence | -0.522 | -1.267 | - | 0.205 | -1.330, 0.290 | 41.073 |
|  |  | Site_category_road | -1.189 | -2.972 | - | **0.003** | **-1.975, -0.400** | 70.938 |
|  |  | Lockdown_status*Site_category_residence | 0.430 | 10.786 | - | **< 0.001** | **0.344, 0.506** | 55.338 |
|  |  | Lockdown_status*Site_category_road | 0.523 | 12.416 | - | **< 0.001** | **0.441, 0.607** | 71.456 |
|  |  | Lockdown_status*Noise | -0.017 | -3.563 | - | **< 0.001** | **-0.026, -0.007** | 1.770 |
|  |  | Lockdown_status*Human_activity | 0.001 | 15.772 | - | **< 0.001** | **0.0006, 0.0008** | 0.106 |
|  | **^#^Activity**  **ariability-**  CV of the  number of  events/day | (Intercept) | -0.316 | - | -2.604 | 0.010 | -0.630, -0.075 | - |
|  |  | Lockdown_status | 0.200 | - | 5.873 | **< 0.001** | **0.047, 0.439** | - |
|  |  | Noise | 0.018 | - | 9.637 | **< 0.001** | **0.014, 0.024** | - |
|  |  | Site_category_residence | 0.087 | - | 1.035 | 0.314 | -0.079, 0.254 | - |
|  |  | Site_category_road | 0.201 | - | 2.467 | **0.023** | **0.034, 0.358** | - |
|  |  | Human_activity | 0.00001 | - | 0.301 | 0.763 | -0.0001, 0.0001 | - |
|  |  | Temperature | -0.008 | - | -2.608 | **0.013** | **-0.013, -0.002** | - |
|  |  | Count_down*Lockdown_status | -0.003 | - | -4.162 | **< 0.001** | **-0.005, -0.002** | - |
|  |  | Lockdown_status*Site_category_residence | -0.050 | - | -1.867 | 0.063 | -0.109, 0.004 | - |
|  |  | Lockdown_status*Site_category_road | -0.104 | - | -3.903 | **< 0.001** | **-0.156, -0.043** | - |
|  |  | Lockdown_status*Human_activity | -0.00001 | - | -2.811 | **0.005** | **-0.0002, -0.00002** | - |
| Graceful  prinia | **Activity-**  Number of  events/day | (Intercept) | 4.013 | 2.593 | - | 0.010 | - | - |
|  |  | Lockdown_status | 1.008 | 1.714 | - | 0.086 | - | 174.012 |
|  |  | Human_activity | 0.0001 | 1.984 | - | 0.047 | - | 0.010 |
|  |  | Noise | -0.198 | -20.677 | - | **< 0.001** | - | 17.963 |
|  |  | Temperature | 0.395 | 6.631 | - | **< 0.001** | - | 48.438 |
|  |  | Site_category_residence | -1.254 | -1.797 | - | 0.072 | - | 72.179 |
|  |  | Site_category_road | -1.424 | -2.093 | - | 0.036 | - | 77.444 |
|  |  | Lockdown_status*Count_down | -0.105 | -15.002 | - | **< 0.001** | - | 10.267 |
|  |  | Lockdown_status*Site_category_residence | -0.041 | -0.744 | - | 0.457 | - | 4.178 |
|  |  | Lockdown_status*Site_category_road | -1.036 | -11.776 | - | **< 0.001** | - | 67.739 |
|  |  | Lockdown_status*Noise | 0.035 | 3.898 | - | **< 0.001** | - | 3.776 |
|  |  | Lockdown_status*Human_activity | 0.0005 | 6.357 | - | **< 0.001** | - | 0.054 |
|  | **^#^Activity**  **ariability-**  CV of the  number of  events/day | (Intercept) | 0.698 | - | 1.548 | 0.123 | -0.472, 1.608 | - |
|  |  | Lockdown_status | -0.324 | - | -2.978 | **0.003** | -0.824, 0.361 | - |
|  |  | Noise | 0.024 | - | 2.903 | **0.004** | **0.006, 0.043** | - |
|  |  | Temperature | -0.040 | - | -3.538 | **0.005** | **-0.059, -0.012** | - |
|  |  | Site_category_residence | 0.211 | - | 0.983 | 0.336 | -0.165, 0.680 | - |
|  |  | Site_category_road | 0.432 | - | 2.036 | 0.052 | **0.078, 0.961** | - |
|  |  | Count_down*Lockdown_status | 0.005 | - | 1.608 | 0.109 | -0.001, 0.011 | - |
|  |  | Lockdown_status*Site_category_residence | 0.146 | - | 1.316 | 0.189 | -0.079, 0.382 | - |
|  |  | Lockdown_status*Site_category_road | 0.285 | - | 2.245 | **0.025** | **0.025, 0.536** | - |

CV: coefficient of variance. *: Interaction effect. 95% confidence intervals of the parameters that did not overlap zero are indicated in bold. **#**: model average.

**Table S19** Akaike’s information criterion (AICc) model comparison results for activity and activity variability in all species and each bird species.

| Species | Model description | *df* | logLik | AICc | ∆AICc | *wi* |
| --- | --- | --- | --- | --- | --- | --- |
| **All species** | **Activity (Number of events/day)** ~ |  |  |  |  |  |
|  | Bird, Lock, Noise, Site, Human, Temperature, Bird*Lock, Count*Lock, Lock*Noise, Lock*Site, Lock*Human | 16 | -29663.59 | 59359.52 | 0.00 | 0.90 |
|  | Bird, Lock, Noise, Site, Human, Temperature, Bird*Lock, Noise*Lock, Lock*Site, Lock*Human | 15 | -29666.80 | 59363.90 | 4.38 | 0.10 |
| **All species** | **Activity Variability (CV of the number of events/day) ~** |  |  |  |  |  |
|  | Bird, Lock, Noise, Site, Human, Temperature, Bird*Lock, Lock*Noise | 13 | 237.72 | -449.20 | 0.00 | 0.23 |
|  | Bird, Lock, Noise, Site, Human, Temperature, Bird*Lock, Count*Lock, Lock*Noise | 14 | 238.72 | -449.15 | 0.05 | 0.22 |
|  | Bird, Lock, Noise, Site, Human, Temperature, Bird*Lock, Lock*Noise, Lock*Human | 14 | 238.35 | -448.41 | 0.79 | 0.16 |
|  | Bird, Lock, Noise, Site, Human, Temperature, Bird*Lock, Count*Lock, Lock*Noise, Lock*Human | 15 | 239.35 | -448.38 | 0.82 | 0.15 |
|  | Bird, Lock, Noise, Site, Temperature, Bird*Lock, Count*Lock, Lock*Noise | 13 | 236.51 | -446.77 | 2.42 | 0.07 |
|  | Bird, Lock, Noise, Site, Human, Temperature, Bird*Lock, Lock*Noise, Lock*Site | 15 | 238.46 | -446.60 | 2.60 | 0.06 |
|  | Bird, Lock, Noise, Site, , Temperature, Bird*Lock, Lock*Noise | 12 | 235.30 | -446.38 | 2.82 | 0.06 |
|  | Bird, Lock, Noise, Site, Human, Temperature, Bird*Lock, Count*Lock, Lock*Noise, Lock*Site | 16 | 239.24 | -446.11 | 3.09 | 0.05 |
| **Hooded crow** | **Activity (Number of events/day)** ~ |  |  |  |  |  |
|  | Lock, Noise, Site, Human, Count*Lock, Lock*Site, Lock*Human | 12 | -5349.02 | 10722.64 | 0.00 | 0.36 |
|  | Lock, Noise, Site, Human, Count*Lock, Lock*Noise, Lock*Site, Lock*Human | 13 | -5348.03 | 10722.76 | 0.12 | 0.34 |
|  | Lock, Noise, Site, Human, Temperature, Count*Lock, Lock*Site, Lock*Human | 13 | -5348.81 | 10724.31 | 1.67 | 0.16 |
|  | Lock, Noise, Site, Human, Temperature, Count*Lock, Lock*Noise, Lock*Site, Lock*Human | 14 | -5347.88 | 10724.56 | 1.92 | 0.14 |
| **Hooded crow** | **Activity Variability (CV of the number of events/day) ~** |  |  |  |  |  |
|  | Lock, Noise, Site, Human, Lock*Noise, Lock*Human | 11 | 245.65 | -468.80 | 0.00 | 0.25 |
|  | Lock, Noise, Site, Human, Temperature, Lock*Noise, Lock*Human | 12 | 246.67 | -468.75 | 0.05 | 0.25 |
|  | Lock, Noise, Site, Human, Count*Lock, Lock*Noise, Lock*Human | 12 | 246.26 | -467.93 | 0.87 | 0.16 |
|  | Lock, Noise, Site, Human, Temperature, Count*Lock, Lock*Noise, Lock*Human | 13 | 246.83 | -466.98 | 1.82 | 0.10 |
|  | Lock, Noise, Site, Human, Lock*Noise, Lock*Site, Lock*Human | 13 | 246.28 | -465.87 | 2.93 | 0.06 |
|  | Lock, Noise, Site, Human, Temperature, Lock*Noise, Lock*Site, Lock*Human | 14 | 247.26 | -465.72 | 3.08 | 0.05 |
|  | Lock, Noise, Site, Human, Count*Lock, Lock*Noise, Lock*Site, Lock*Human | 14 | 247.15 | -465.49 | 3.31 | 0.05 |
|  | Lock, Noise, Human, Lock*Noise, Lock*Human | 9 | 241.62 | -464.89 | 3.91 | 0.04 |
|  | Lock, Noise, Human, Temperature, Lock*Noise, Lock*Human | 10 | 242.63 | -464.85 | 3.95 | 0.04 |
| **Rose-ringed**  **parakeet** | **Activity (Number of events/day)** ~ |  |  |  |  |  |
|  | Lock, Noise, Site, Human, Temperature, Lock*Noise, Lock*Site, Lock*Human | 13 | -3599.12 | 7224.94 | 0.00 | 0.59 |
|  | Lock, Noise, Site, Human, Temperature, Count*Lock, Lock*Noise, Lock*Site, Lock*Human | 14 | -3598.44 | 7225.69 | 0.75 | 0.41 |
| **Rose-ringed**  **parakeet** | **Activity Variability (CV of the number of events/day) ~** |  |  |  |  |  |
|  | Lock, Noise, Site, Human, Temperature, Count*Lock, Lock*Site, Lock*Human | 14 | 460.70 | -892.59 | 0.00 | 0.65 |
|  | Lock, Noise, Site, Human, Temperature, Count*Lock, Lock*Noise, Lock*Site, Lock*Human | 15 | 461.16 | -891.39 | 1.20 | 0.35 |
| **Graceful prinia** | **Activity (Number of events/day)** ~ |  |  |  |  |  |
|  | Lock, Noise, Site, Human, Temperature, Count*Lock, Lock*Noise, Lock*Site, Lock*Human | 14 | -7795.18 | 15619.17 | 0.00 | 1.00 |
|  | Lock, Noise, Site, Human, Temperature, Count*Lock, Lock*Site, Lock*Human | 13 | -7802.84 | 15632.38 | 13.21 | 0.00 |
| **Graceful prinia** | **Activity Variability (CV of the number of events/day) ~** |  |  |  |  |  |
|  | Lock, Noise, Site, Temperature, Count*Lock, Lock*Site, Lock*Human | 12 | -225.14 | 475.06 | 0.00 | 0.16 |
|  | Lock, Noise, Site, Temperature, Lock*Site | 11 | -226.43 | 475.52 | 0.45 | 0.12 |
|  | Lock, Noise, Site, Temperature | 9 | -228.62 | 475.69 | 0.63 | 0.11 |
|  | Lock, Noise, Site, Temperature, Count*Lock | 10 | -227.65 | 475.85 | 0.78 | 0.11 |
|  | Noise, Site, Temperature | 8 | -230.19 | 476.74 | 1.68 | 0.07 |
|  | Lock, Noise, Site, Human, Temperature, Count*Lock, Lock*Site | 13 | -225.02 | 476.97 | 1.90 | 0.06 |
|  | Lock, Noise, Site, Temperature, Count*Lock, Lock*Noise, Lock*Site | 13 | -225.12 | 477.17 | 2.10 | 0.05 |
|  | Lock, Noise, Site, Human, Temperature, Lock*Site | 12 | -226.36 | 477.51 | 2.45 | 0.05 |
|  | Lock, Noise, Site, Temperature, Lock*Noise | 10 | -228.53 | 477.62 | 2.55 | 0.04 |
|  | Lock, Noise, Site, Temperature, Lock*Noise, Lock*Site | 12 | -226.42 | 477.63 | 2.57 | 0.04 |
|  | Lock, Noise, Site, Human, Temperature | 10 | -228.62 | 477.80 | 2.73 | 0.04 |
|  | Lock, Noise, Site, Temperature, Count*Lock, Lock*Noise | 11 | -227.59 | 477.85 | 2.78 | 0.04 |
|  | Lock, Noise, Site, Human, Temperature, Count*Lock | 11 | -227.64 | 477.94 | 2.88 | 0.04 |
|  | Noise, Site, Human, Temperature | 9 | -230.19 | 478.83 | 3.77 | 0.02 |
|  | Lock, Noise, Site, Human, Temperature, Count*Lock, Lock*Site, Lock*Human | 14 | -224.92 | 478.91 | 3.85 | 0.02 |
|  | Lock, Noise, Site, Human, Temperature, Count*Lock, Lock*Noise, Lock*Site | 14 | -225.00 | 479.06 | 3.99 | 0.02 |

Bird: Bird species. Lock: Lockdown status. Site: Site category. Human: Human activity. Count: Count down. AICc: Akaike’s information criterion corrected for small sample size. ΔAICc: the difference between the alternative model and best-fitting model. Models are ranked based on the AICc values from the best to the worst model. *: Interaction effect.
